# Correlation between speed and turning naturally arises for sparsely sampled cell movements

**DOI:** 10.1101/2020.12.30.424897

**Authors:** Vitaly V. Ganusov, Viktor S. Zenkov, Barun Majumder

**Affiliations:** Department of Microbiology, University of Tennessee, Knoxville, TN 37996, USA; Department of Mathematics, University of Tennessee, Knoxville, TN 37996, USA; Department of Electrical Engineering and Computer Science, University of Tennessee, Knoxville, TN 37996, USA

## Abstract

Mechanisms regulating cell movement are not fully understood. One feature of cell movement that determines how far cells displace from an initial position is persistence, the ability to perform movements in a direction similar to the previous movement direction. Persistence is thus determined by turning angles between two sequential displacements and be characterized by an average turning angle or persistence time. Recent studies found that a cell’s average speed and turning are negatively correlated, suggesting a fundamental cell-intrinsic program whereby cells with a lower turning ability (i.e., larger persistence time) are intrinsically faster (or faster cells turn less). By simulating correlated or persistent random walks (PRWs) using two different frameworks (one based on von Mises-Fisher (vMF) distribution and another based on Ornstein-Uhlenbeck (OU) process) we show that the negative correlation between speed and turning naturally arises when cell trajectories are sub-sampled, i.e., when the frequency of sampling is lower than frequency at which cells make movements. This effect is strongest when the sampling frequency is on the order of magnitude with the typical cell persistence time and when cells vary in persistence time. Both conditions are observed for datasets of T cell movements in vivo that we have analyzed. In simulations the correlation arises due to randomness of cell movements resulting in highly variable persistence times for individual cells that with sub-sampling leads to large variability of average cell speeds. Interestingly, previously suggested methodology of calculating displacement of cell cohorts with different speeds resulted in similar results whether or not there is a cell-intrinsic correlation between cell speed and persistence. For both vMF- and OU-based simulations of PRWs we could find parameter values (distribution of persistence times, speeds, and sampling frequency) that matched experimentally measured correlations between speed and turning for two datasets of T cell movement in vivo suggesting that such simple correlations are not fully informative on the intrinsic link between speed and persistence. Our results thus suggest that sub-sampling may contribute to (and perhaps fully explains) the observed correlation between speed and turning at least for some cell trajectory data and emphasize the role of sampling frequency in inference of critical cellular parameters of cell motility such as speeds.

**Secondary Abstract:** Measurement of cell movements often results in a negative correlation between average speed and average turning angle suggesting an existence of a universal, cell-intrinsic movement program. We show that such a negative correlation may arise if cells in the population differ in their ability for persistent movement when the movement data are sub-sampled. We show that sub-sampling of cell trajectories generated using two different frameworks of persistent random walk can match the experimentally observed correlation between speed and turning for T cell movements in vivo.

## Introduction

Motility is a fundamental property of cells. Motility is exhibited by single cell organisms such as bacteria or ameba as well as by cells of multicellular organisms such as cells during development or cancer [1–4]. To protect the host against pathogens, T lymphocytes, cells of the adaptive immune system, need to constantly move and survey tissues for infection [5, 6]. While molecular mechanisms of T cell motility have been relatively well defined, specific strategies that T cells use to efficiently locate the pathogen remain controversial. Some studies indicated that T cells search for the infection in the lymph nodes randomly [7, 8] while others suggested an important role of attracting cues such as chemokines [6, 9]. In non-lymphoid tissues, a wider range of movement types of T cells have been observed, including Brownian walks in the skin [10], correlated random walks in the liver [11] or in explanted lungs [12], or generalized Levy walks in murine brains [13].

Movement of cells in tissues *in vivo* is typically recorded using intravital microscopy at a particular frame rate (e.g., a small 3D volume of the liver or a lymph node of 500 × 500 × 50 *µ*m can be scanned with a two photon microscope every 20-30 sec, [11, 14, 15]). By segmenting images with software packages (e.g., ImageJ from the NIH or Imaris from Bitplane), 3D coordinates of multiple cells over time can be obtained. Generation of 3D cell coordinates from raw imaging data is a complicated process, and in some cases *z*-coordinates are either ignored or collapsed into one using maximal projection, resulting only in change in *x* and *y* coordinates for individual cells over time [16–18]. Cell coordinates can be then used to characterize movement pattern of the cells. Several alternative parameters are useful in this regard [17, 19]. One is the distribution of movement lengths, which when adjusted based on the imaging frequency gives the distribution of instantaneous speeds of cells from which the average speed per cell can be calculated. The distribution of movement lengths can be used to infer heterogeneity in cell migration speeds or if cells are utilizing a specific search strategy such as Levy walks [11, 13, 16, 20, 21]. It is important to note, however, that the estimated average or instantaneous speeds of cells are generally dependent on the frequency at which imaging is performed [22].

Another important parameter characterizing cell movement is persistence - the ability of cells to keep moving in the direction similar to the previous movement direction; such a walk type is also defined as a correlated or persistent random walk (PRW). While intuitively cell persistence is a relatively clear concept, ways to characterize how “persistent” cells in a given population are, vary. One approach is to perform linear regression analysis on the initial change in the log-transformed mean square displacement (MSD) of cells vs. log-transformed time and find the time *T*_*p*_ until which the slope in this initial change is larger than 1. A more rigorous approach is to fit the Fürth equation to the MSD curve and estimate persistence time as one of the equation’s parameters [16, 18, 23–25]. Another way is to calculate the slope in the exponential decline of the correlation between cell’s velocity vectors over time when averaged over all cells in the population; an inverse value of the slope also gives the average persistence time [17, 18, 24, 25]. An alternative approach to characterize the ability of cells in the population to undergo a correlated random walk is to calculate turning angles - angles between consecutive movement vectors in an experimental movie [19, 22, **Figure 1**]. The distribution of turning angles along with the movement length distribution, thus, characterizes the movement pattern of cells in the population. The average turning angle 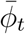 may then inform the ability of cells in the population to persistently move: when the average turning angle is 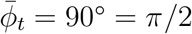 (and the turning angle distribution is described by a sin function), cells are not persistent. The fraction of cell movements with turning angles higher than 90^*°*^ also indicates the probability of cells to turn away from the previous movement direction. However, as compared to the estimated persistent time *T*_*p*_, the average turning angle 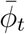 does not represent an intuitive parameter indicating how long cells are persistent in their movement. Yet, while calculating the average turning angle for individual cells is possible for datasets of different sizes, calculating persistence times for individual cells can be problematic for datasets of cell movement *in vivo* since such data typically contain *<* 100 movements per cell (e.g., [11, 13]).

**Figure 1:**
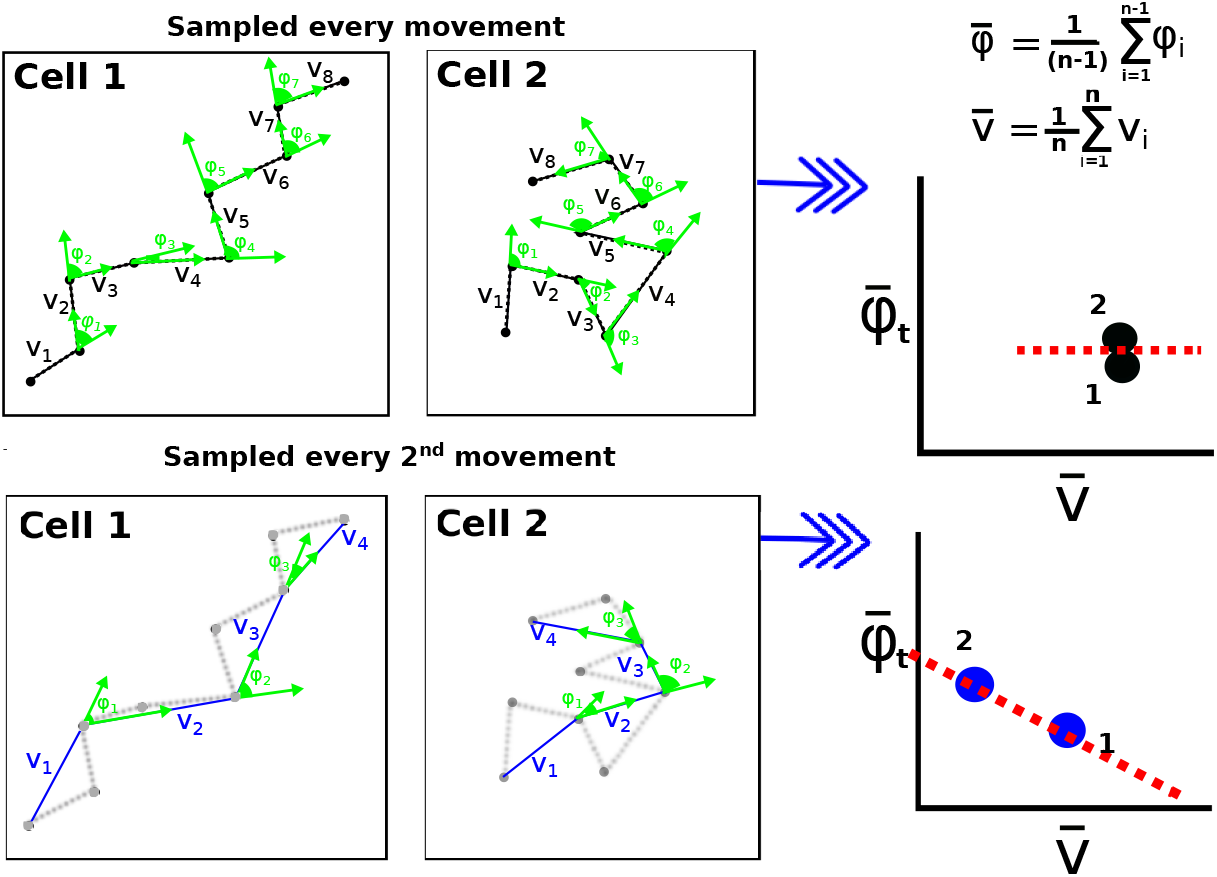
Schematic representation of how frequency of sampling of cell movement influences the observed relationship between average speed and average turning angle per cell. We plot trajectories for 2 cells and show the speed of every movement (denoted as *v*_*i*_) and turning angle 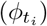. When sampling is done at the same frequency as cells make a decision to turn (top panels), per model assumption average speed 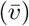 and average turning angle 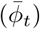 do not correlate. However, when sampling is done less frequently (e.g., every 2nd movement, bottom panels) the cell 2 that turned more has a lower average speed and higher average turning angle as compared to the cell 1, which turned less and had a more persistent walk. High sensitivity of the estimated average speed to the frequency at which movements are sampled is important in generating the observed correlation between speed and persistence.

What determines the ability of cells to exhibit correlated random walks remains poorly understood. We recently argued that the constrained environment of the liver sinusoids is likely to force liver-localized CD8 T cells to perform correlated random walks resulting in super-diffusive displacement for 10-15 minutes [11]. However, the ability of cells to perform a correlated random walk could be cell-intrinsic. Indeed, authors of several recent studies accurately measured movement of cells in 2D (*in vitro*) or in 3D (*in vivo*) over time and found that there is a strong positive correlation between average cell persistence, defined as the persistence time or the cosine of turning angles, and average cell speed [18, 26, 27]; this is equivalent to the negative correlation between average turning angle per cell and cell speed. Specifically, experiments by Jerison & Quake [18] were designed to monitor migration of T cells *in vivo* in zebrafish for hours given the transparency of the animal and methods to stitch different images across nearly the whole animal. Importantly, T cell tracking was performed in 2D with a single-plane illumination microscope [18]. Their analysis of the data suggested that T cells exhibited large heterogeneity in movement speeds, cohorts of cells with similar speeds showed different persistence times, and there was a strong negative correlation between average turning angle per cell and average speed [18, see also Results section], suggesting a fundamental property of cell movement: cells that turn less have intrinsically larger speeds (or that faster cells turn less) [18, 26]. Importantly, the correlation between turning ability and speed, sometimes called Universal Coupling between Speed and Persistence (UCSP), was observed for different cell types, including unicellular organisms, and genetic perturbations or treatment of cells with drugs impacting cell movement ability did not eliminate the observed correlation, suggesting that indeed the relationship between persistence and speed may be fundamental to cell movement [18, 26, 28].

It is important to recognize that both cell persistence (evaluated, say, by average turning angle) and cell speed are estimated parameters from experimental data, and the true turning ability and true instantaneous speeds of cells are not generally known. The frequency at which individual cells make decisions to change direction of movement and movement speed is also unknown, especially given imprecise measurements of cell positions at a high frequency of imaging *in vivo*. Here we show that the experimentally observed negative correlation between average turning angle and average speed per cell naturally arises in simulations of cells undergoing correlated/persistent random walks due to coarse sampling of cell movement trajectories (**Figure 1**). Indeed, when cells undergo a correlated random walk some cells may displace far and some may stay localized due to random chance alone, and when trajectories are sub-sampled, one expects to see a negative correlation between average turning angle and average speed (**Figure 1**) because cells which remain localized tend to show lower measured speed with large turns compared to the ones which displace far. We show that when there is a variability in persistence ability between individual cells (but not a variability in speed), the negative correlation between average turning angle and average speed is observed for a large range of measured speeds. We found sets of parameters for distribution of persistence and speeds that allow to relatively well match correlations between average speed and average turning angle found in two independent experimental datasets of T cell movement *in vivo*. Importantly, none of the conventionally used parameters such as MSD for cell cohorts or the slope between average turning angle and average speed allowed us to discriminate between the two hypotheses in which there is or there is not a cell-intrinsic link between persistence and speed. We also showed that the negative correlation between average turning angle and speed strongly depends on the relative ratio of the imaging frequency and typical cell persistence time. Our results, thus, suggest that sub-sampling of cell movements may in some cases contribute to the observed correlation between speed and turning.

## Results

### Correlation between persistence and speed arises naturally in simulations of PRWs with sub-sampled data

Because cells’ intrinsic programs for speed and ability to turn are not known, a cell’s speed and average turning angle are estimated using measurements. Geometrically, it is clear that if the sampling of a cell’s positions occurs rarer than the times of decisions the cell makes to turn, a negative correlation between the estimated speed and estimated turning angle may naturally arise because cells that turned less by chance are likely to be observed to displace further, and thus have a higher estimated speed (**Figure 1**). Likewise, the cells that remain localized will have the same displacement but the observed speed will be low because of sampling at larger time steps. This logic remains valid if we were to measure cell’s persistence with other metrics that may be less dependent on sampling frequency (e.g., persistence time, see below) because inferred speed is still sensitive to sampling frequency. To check this intuition, we ran a series of stochastic simulations.

First, we simulated 500 cells moving randomly via Brownian motion with movement lengths following a thin-tailed distribution (Pareto distribution with *α* = 5.5 and 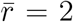, [11, see Materials and Methods]). When the cells’ positions were sampled at the frequency that the cells were changing movement direction (i.e., *k* = 1), we observed no correlation between average speed and average turning angle as assumed (**Figure 2**A and **Supplemental Figure S1**). As expected with sampling every *k* = 10^th^ movement, we still observed no correlation between average speed and turning angle (**Figure 2**B). The average speed of cells in such rarely sampled data was also about 4 times smaller than the assumed speed, because cells were turning and thus not displacing far from the initial location.

**Figure 2:**
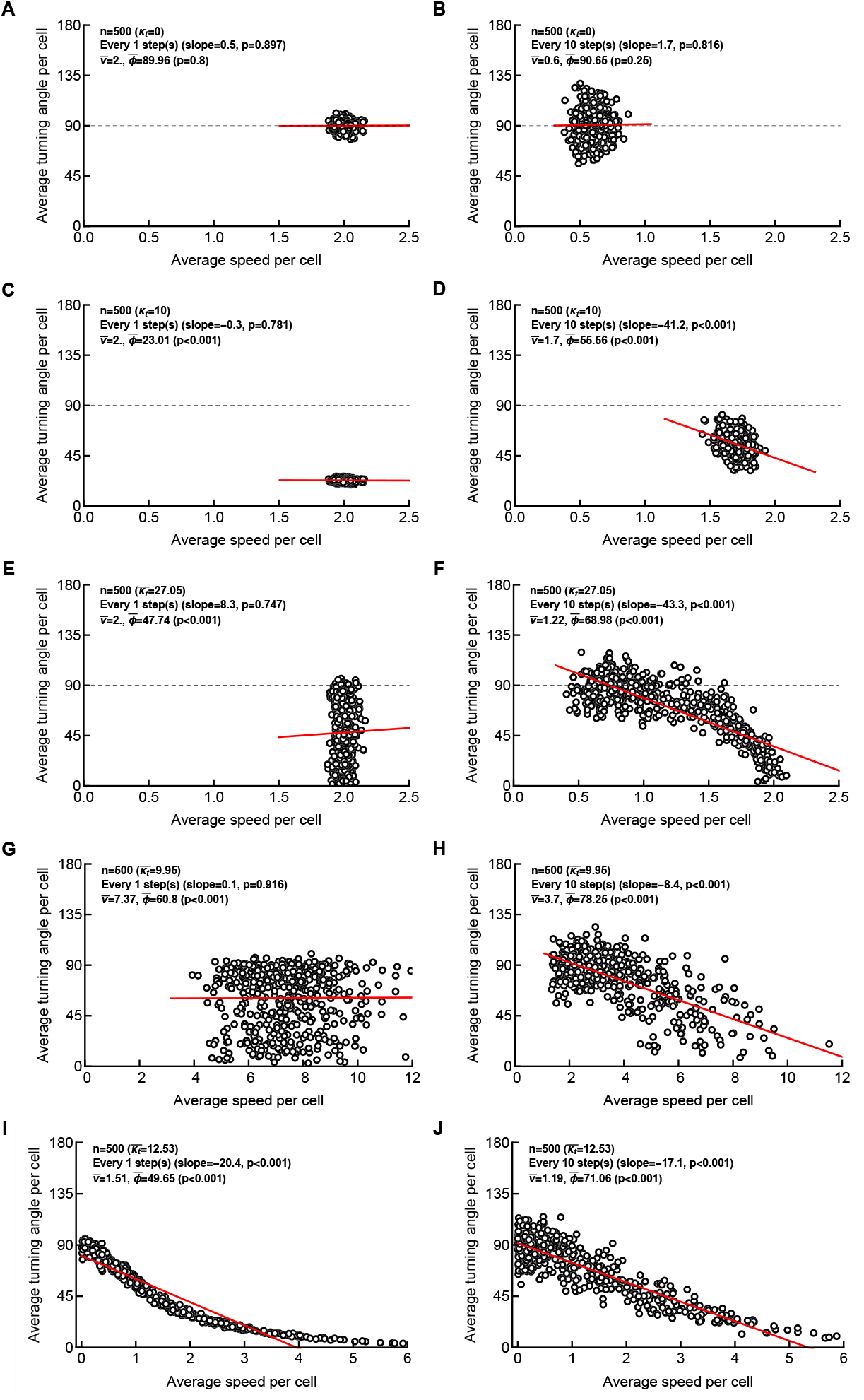
Correlation between average speed and average turning angle may arise in the absence of a cell-intrinsic link between cells’ speed and cells’ turning angles due to sub-sampling. We simulated movement of 500 cells using vMF distribution assuming i) Brownian walk (*κ*_*t*_ *→* 0, A-B), ii) persistence for forward movement being identical for all cells (*κ*_*t*_ = 10, C-D), iii) heterogeneity in cells’ persistence of movement (*κ*_*t*_ was sampled from a lognormal distribution with *µ* = 0.2 and *σ* = 2, E-F), iv) independent heterogeneity in cells’ persistence and speed movement (*κ*_*t*_ and 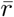 were sampled from a lognormal distribution with *µ* = 0 and *σ* = 2 for *κ*_*t*_ and with *µ* = 2 and *σ* = 0.2 for 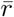, v) a direct relationship between cells’ persistence ability defined by *κ*_*t*_ and cells’ intrinsic movement speed (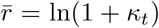), with *κ*_*t*_ following a lognormal distribution with *µ* = 1 and *σ* = 2, I-J). The resulting trajectories were sampled either every step (A, C, E, G, I) or every *k* = 10^th^ step (B, D, F, H, J). Other details of simulations are given in Materials and Methods. Each panel contains information on the average speed for all cells 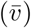, average turning angle for all cells 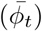, and the result of linear regression of the average speed per cell and average turning angle per cell (denoted as “slope” and shown by red line) with *p* value from the t-test. We also test if the average turning angle of cells in the population is different from 90^*°*^ (Mann-Whitney test). Note different scales in A-F and G-J due to higher speeds of cells in simulations in G-J.

Second, to simulate a correlated random walk we used our newly proposed methodology of sampling random vectors with a bias towards a particular direction using the von Mises-Fisher (vMF) distribution [29, see Materials and Methods]. When we allowed for a moderately biased correlated random walk (with the concentration parameter *κ*_*t*_ = 1 in the vMF distribution corresponding approximately to the persistence time of 1 step, see below), we did not observe a statistically significant correlation between average turning angle and average speed in sub-sampled data. However, for a more biased correlated random walk (*κ*_*t*_ = 10), we observed that rarer (*k* = 10) but not regular (*k* = 1) sampling of cells’ positions resulted in a statistically significant but weak negative correlation between measured speed and the average turning angle per individual cell (**Figure 2**C-D and **Supplemental Figure S2**). This result arose because even with the assumed identical parameter for cell persistence *κ*_*t*_ due to randomness, some cells exhibited walks with long persistence while other cells turned (**Supplemental Figure S3**) resulting in a distribution of persistence times between individual cells. Importantly, sub-sampling of every 3 steps (*k* = 3) already resulted in a statistically significant correlation between speed and turning suggesting that even small sub-sampling can generate a spurious correlation (**Supplemental Figure S2**C).

However, it is possible that cells differ in their ability to turn, e.g., either because of cell-intrinsic program or because of environmental constraints by physical barriers or chemical cues in specific locations of the tissue [11]. Therefore, in the third set of simulations we allowed every cell to have an intrinsic turning ability (defined by individual *κ*_*t*_) drawn from a lognormal distribution (to allow for a broader range of *κ*_*t*_). Importantly, while there was no correlation between speed and turning angle for frequently measured cell movements (**Figure 2**E), when sub-sampling the trajectory data we observed a strong negative correlation between speed and turning angle for a larger span of the speeds (**Figure 2**F and **Supplemental Figure S4**). Impressively, sub-sampling only other step (*k* = 2) already resulted in statistically significant correlation between speed and turning (**Supplemental Figure S4**C). Similar to “simpler” simulations, cells that had a smaller *κ*_*t*_ had a higher propensity to turn, resulting in a smaller overall displacement, and thus, in smaller measured speeds. In contrast, cells that turn little (high *κ*_*t*_) have estimated speeds close to the set intrinsic value 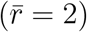. Interestingly, allowing for speeds to be a cell’s property (i.e., when 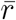 for individual cells was sampled from a lognormal distribution) with cells undergoing a persistent random walk (*κ*_*t*_ = 10) did not result in a negative correlation between speed and turning angle suggesting that the relative variability in walk persistence (determined by *κ*_*t*_) and in speeds (determined by 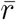) are important for the observed correlation in sub-sampled data (**Supplemental Figure S5**).

In the fourth set of simulations, we allowed both speed 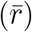 and persistence (*κ*_*t*_) to vary between individual cells independently. Using a lognormal distribution allowed for a range of speeds and average turning angles which were independent for frequently sampled data (**Figure 2**G), but there was a strong negative correlation between average turning angle and average speed for sub-sampled trajectories (**Figure 2**H and **Supplemental Figure S6**) which resembled experimentally observed correlations (see below). Again, sub-sampling every other step (*k* = 2) already resulted in negative correlation between speed and turning (**Supplemental Figure S6**D).

Fifth and finally, we tested how the frequency of sampling of cell movements influences the observed correlation between cell speed and average turning angle when there is an intrinsic link between the instantaneous speed of the cell and its turning ability. We therefore simulated cell movement by sampling *κ*_*t*_ from a lognormal distribution and then linked cells’ intrinsic speed to cells’ turning ability (by assuming 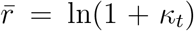 as one example). Interestingly, the frequency of sampling had a moderate effect on the negative correlation between average speed and average turning angle (**Figure 2**I-J and **Supplemental Figure S7**). Taken together, these results strongly suggest that because the intrinsic cell speed, intrinsic turning ability, or frequency at which any cell makes decisions of turning are not known, a negative correlation between measured speed of cells and average turning angle may arise due to sub-sampling.

### Correlation between average turning angle and speed is observed only for a range of sampling frequencies

The frequency of sampling of cell trajectories relative to a typical persistence time of cells in the population should impact the correlation between speed and turning. In particular, if sampling is too coarse and exceeds the persistence time for all cells there should be no correlation between average turning angle and speed. Similarly, if the sampling occurs at the frequency at which cells make movements, turning and speed should not be correlated (if there is no intrinsic link between speed and persistence).

To check this intuition we performed a set of longer simulations of PRWs with vMF distribution. Specifically, we simulated 300 cells traversing for 1000 steps assuming that either all the cells have same ability of persistence (defined by *κ*_*t*_ of the vMF distribution) and same speed (defined by 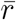) or when there is a distribution of *κ*_*t*_ and 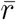 between individual cells. Importantly, in both cases when we sample every cell movement, the correlation between average turning angle and speed is not statistically significant, while at some intermediate values of the sampling frequency *k* it becomes highly significant, but then disappears when sampling is too coarse (**Figure 3**A-B). Importantly, a very similar pattern is observed when speed and turning are linked, i.e., when we assume a lognormal distribution of turning abilities of the cells with speeds being directly determined by *κ*_*t*_ (**Figure 3**C-D).

**Figure 3:**
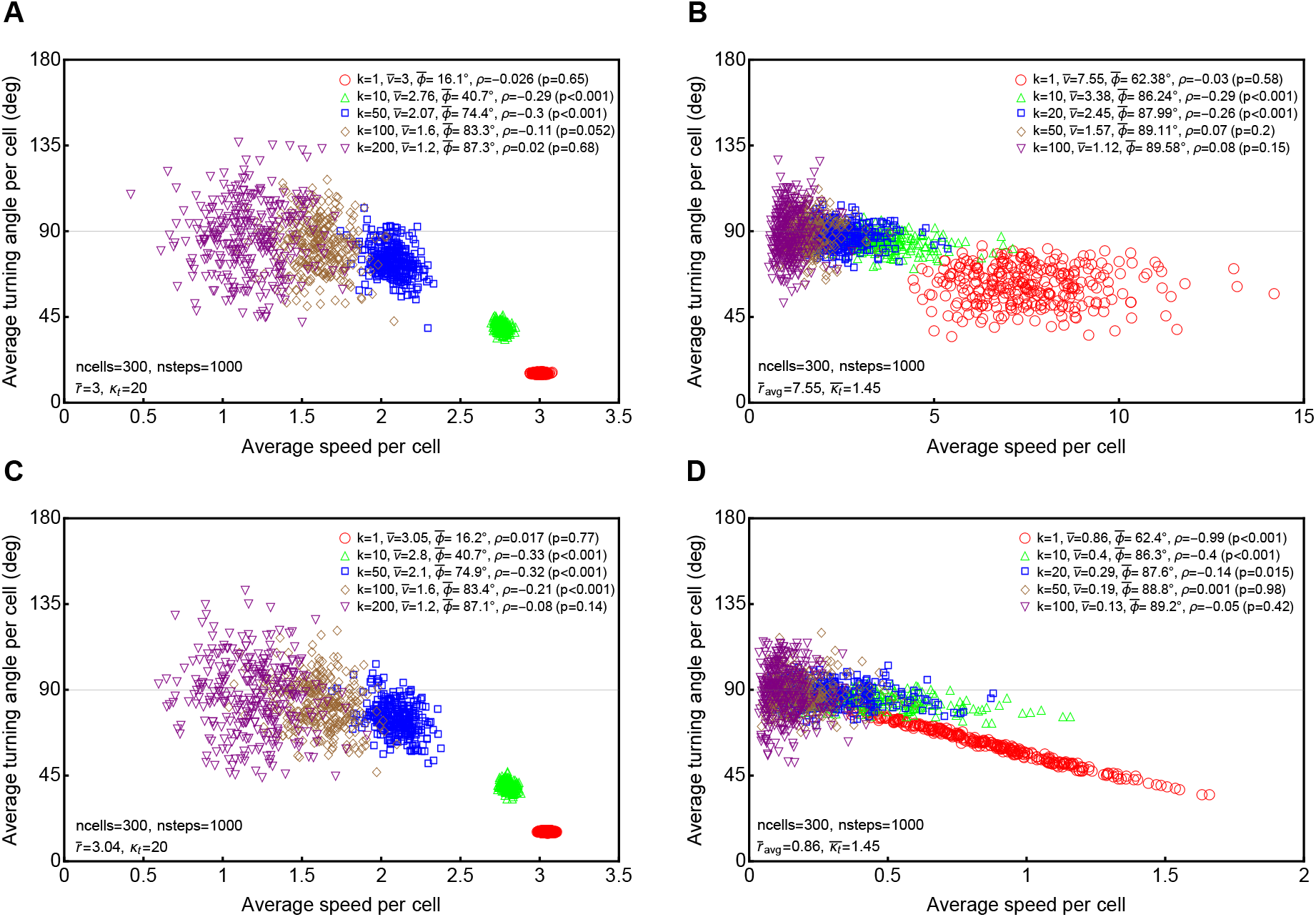
Correlation between speed and turning angle disappears for coarsely sub-sampled simulation data. We simulated 300 cells each with 10^3^ steps using vMF distribution and sampled every *k*^th^ movements (*k* is indicated on individual panels). In panels A-B we assume that speed and concentration parameter *κ*_*t*_ are uncorrelated, and in panels C-D, speed is determined by *κ*_*t*_ via 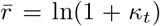. In panels A and C we assume that every cell have the same persistence defined by *κ*_*t*_ = 20 and same speed defined by 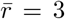. In panels B and D we assume that every cell in the population has a different *κ*_*t*_ which was drawn from a lognormal distribution (eqn. (4) with *µ* = 0.2 and *σ* = 0.5). In panel B, every cell has a random speed determined by 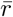 in the Pareto distribution (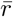 was drawn from a lognormal distribution with *µ* = 2 and *σ* = 0.2), and in panel D speeds are directly determined by *κ*_*t*_ as 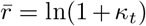. Average speed 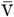 and average turning angle 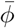 for all cells are indicated on the panels, and statistical significance of the correlation between speed and turning angle per cell was determined using Spearman rank test (with the correlation coefficient *ρ* and p-values are shown on individual panels).

We performed another set of simulations where we varied the average persistence per cell (defined by *κ*_*t*_) but fixed the sampling frequency of the trajectories (*k* = 20). Interestingly, we found a statistically significant correlation between average speed and average turning angle per cell for a broad range of *κ*_*t*_ (*κ*_*t*_ = 4 − 100, **Supplemental Figure S8**). This result suggests that the correlation between speed and turning is most likely to arise when the sampling frequency of cell movements is of the same order of magnitude as a typical persistence time of cells in the population (*k ∼ κ*_*t*_).

### Average turning angle and persistence time are correlated metrics of cell persistence

In addition to average turning angle, persistence time is another metric that has been used to measure ability of cells to persist. Two methods have been proposed to estimate persistence time following the Ornstein-Uhlenbeck formulation of the PRW. One is by fitting the Fürth equation to the MSD data and another is by fitting an exponential decay function to the velocity correlations [16, 30, and see Materials and Methods for more detail]. We performed simulations of 500 cell tracks, varied the concentration parameter *κ*_*t*_ of the vMF distribution, and estimated persistence times for individual cells (**Supplemental Figure S9**). We found a strong correlation between estimates found by the two methods, although the method based on the decay of velocity correlations provided lower estimates of the persistence time for larger times (**Supplemental Figure S9**).

In our simulations with the vMF distribution the concentration parameter *κ*_*t*_ quantifies the persistence ability of a cell; however, it was not clear how the magnitude of *κ*_*t*_ relates to cell’s persistence time. Therefore, we simulated movement of 500 cells each with 100 steps by fixing the cell’s intrinsic speed 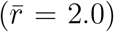 and *κ*_*t*_ for each cell by allowing *κ*_*t*_ to vary between different simulations. We then either sampled every cell position (*k* = 1) or every 10^th^ (*k* = 10) position and estimated the average turning angle 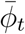 and average persistence time *T*_*p*_ for the population of cells using Fürth equation (eqn. (7)). As expected we found strong positive correlation between *κ*_*t*_ and *T*_*p*_, while average turning angle was a declining function of *κ*_*t*_ (**Supplemental Figure S10**A&B). Also, these results showed a strong negative correlation between persistence time and the turning angle, suggesting that both metrics can be used to evaluate the ability of cells to persist (**Supplemental Figure S10**C). Interestingly, sampling frequency determined by *k* had little influence on the estimated persistence time (**Supplemental Figure S10**A). However, when persistence is small (e.g., *κ*_*t*_ *≈* 1), the Fürth equation did not allow to estimate the persistence time (estimated time was less than 1 step and thus unreliable), and yet, the average turning angle was significantly lower than 90^0^, suggesting a higher sensitivity of the latter metric to detect deviation from random turning angles. Persistence time may be difficult to estimate for individual cells when amount of data available per cell is limited. For example, plots of the velocity correlations for individual cells with identical assumed persistence (*κ*_*t*_ = 10) show large variability that translates into variable persistence time while average turning angle per cell is relatively stable (**Supplemental Figure S11**).

Despite these potential shortcomings of using persistence time as a metric to measure movement persistence of individual cells we investigated whether sub-sampling influences the correlation between persistence time and speed. Therefore, we performed identical simulations to those in **Figure 2**E-H and estimated persistence time of individual cells using velocity correlations. We found that that sub-sampling (*k* = 5) resulted in a positive correlation between persistence time and average speed per cell (**Supplemental Figure S12**B&D) which was absent when all data were used (*k* = 1, **Supplemental Figure S12**A&C). This is because while persistence time of a given cell is relatively insensitivity to sampling frequency, the average speed per cell is not, and sub-sampling results in lower speed estimates for cells with smaller persistence times. Thus, sub-sampling results in correlation between speed and persistence independently of the metric (persistence time or average turning angle) used to quantify cell persistence.

### Correlation between speed and turning arises in another framework to simulate PRWs

All our results so far were found using a novel method of simulating PRWs using the vMF distribution [29]. To check that the correlation between average turning angle and average speed arising due to sub-sampling is model-independent (as logic suggests, e.g., **Figure 1**), we simulated PRWs using an algorithm outlined by Wu *et al*. [16] that is a direct implementation of the Ornstein-Uhlenbeck model of cell movement [23–25]. We simulated 300 cells traversing 7200 steps which is equivalent to 2 hours of movement assuming that each step is 1 sec (see Materials and methods for more detail). We also assumed that either cells in the population have the same persistence time of *T*_*p*_ = 0.75 min and same speed *s* = 4.5 *µ*m/min (**Supplemental Figure S13**A) or that both parameters vary between cells in accord with lognormal distributions (**Supplemental Figure S13**B). Interestingly, we found a weak negative correlation between speed and turning angle in these simulations even when we sample every cell movement (*k* = 1 sec, **Supplemental Figure S13**A). As we sub-sample the data, the correlation between average speed and average turning angle becomes noisier (and more resembles experimental data, see below), but at higher *k*, when the sub-sampling becomes coarser than the typical persistence time, the correlation disappears (**Supplemental Figure S13**). Thus, our results of simulations based on vMF distribution are confirmed with another method to simulate PRWs.

### Several measures such as average turning angle, speed, or MSD for cell cohorts do not allow to discriminate between alternative hypotheses

In our simulations it was clear that the frequency of sampling had a major impact on the regression slope between average cell speed and average turning angle (e.g., **Supplemental Figures S2 and S4**). We therefore tested if a change in the frequency of sampling of cell movement can be used to discriminate between cell-intrinsic vs. randomly arising negative correlation between speed and turning angle. We calculated how the slopes between speed and turning angle, average speed, or average turning angle change when the frequency of imaging *k* changes (**Figure 4**). Unfortunately, the two models were similar in how these parameters changed with imaging frequency, except for the narrow range when imaging frequency would coincide with the frequency at which cells make turn decisions (**Figure 4**A at *k* = 1). However, very frequent imaging contains many artifacts due to changes in cell shape or fluorescence signal from the cell [31] suggesting that changes in the frequency of imaging may not be the method to discriminate between these alternatives.

**Figure 4:**
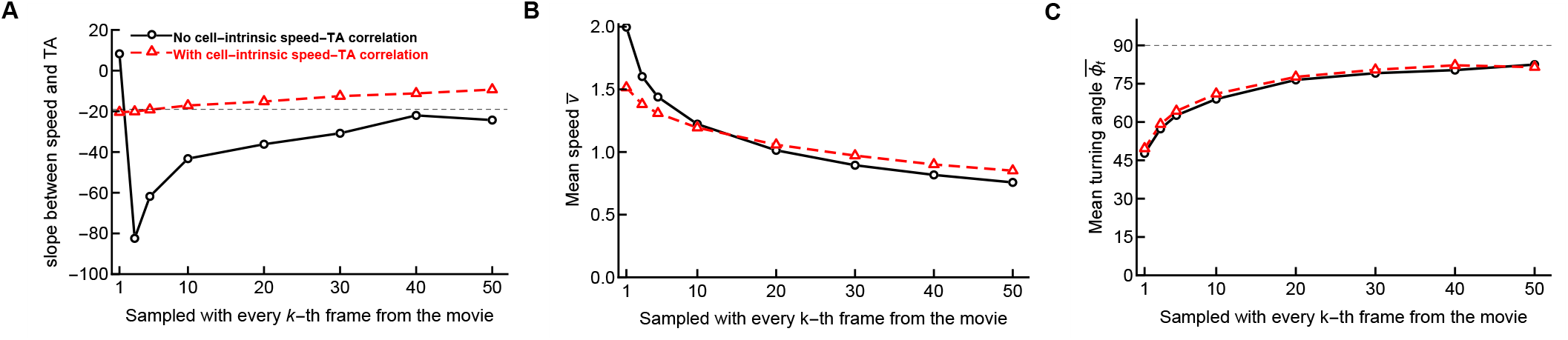
Frequency of imaging does not allow discrimination between hypotheses explaining negative correlation between measured average speed and average turning angle (TA) for a cell. We simulated movement of cells assuming that i) each cell in the population has a preference toward moving forward (defined by the concentration parameter *κ*_*t*_) but all cells have the same intrinsic speed (**Figure 2**E-F) or ii) for each cell is characterized by a preference to move forward defined by *κ*_*t*_ and intrinsic speed defined as 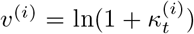 where (*i*) indicates an *i*^th^ cell (**Figure 2**I-J). For different frequencies of recording cell movement, we calculated the slope between average speed and average turning angle (TA or 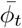) per cell (panel A), the average speed per cell (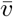, panel B), and the average turning angle per cell (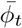, panel C) for the two hypotheses (without and with cell-intrinsic link between speed and turning angle, shown by different markers and lines). The thin dashed line in panel A denotes the expected slope for the model with a cell-intrinsic link between speed and turning at the lowest possible frequency of imaging (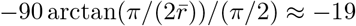 for 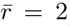). Values on the x-axes indicate which frames were included in the calculations. For example, *k* = 10 indicates that data at *t* = 1, 11, 21, 31 … were used in calculations. In simulations, the concentration parameter *κ*_*t*_ was lognormally distributed between cells with *µ* = 1 and *σ* = 2 (see **Figure 2** and Materials and Methods for more detail).

xThe MSD curves have been also used to establish the linear relationship between persistence and speed for cohorts of cells that have similar speeds [18]. Specifically, cell cohorts with higher speeds had a faster increase in MSD with time (higher *γ*, see eqn. (6) and other details in Materials and methods) that was interpreted as evidence of the direct relationship between cell speed and persistence [18]. We therefore performed additional analyses to determine if we can reject the null model in which turning ability and speeds are cell-dependent but are uncorrelated using this methodology. For this we simulated movements of 500 cells assuming correlated random walks using the vMF distribution. In one set of simulations, we considered the persistence *κ*_*t*_ and the speed characterizing parameter 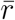, drawn from two independent lognormal distributions (**Figure 5**A-B and **Supplemental Figure S14**). As expected, the frequency of sampling impacted dramatically how different cell cohorts displaced over time (**Supplemental Figure S14**), and while for frequent (every *k* = 1 frame) sampling different cell cohorts were similarly super-diffusive with *γ >* 1 (**Figure 5**A and **Supplemental Figure S14**A), coarse sampling (every *k* = 10 movements) resulted in different cohorts displacing differently, with slowest cohorts having low (*γ* = 1.2, **Figure 5**B) or close to Brownian diffusion (*γ ≈* 1, **Supplemental Figure S14**D).

**Figure 5:**
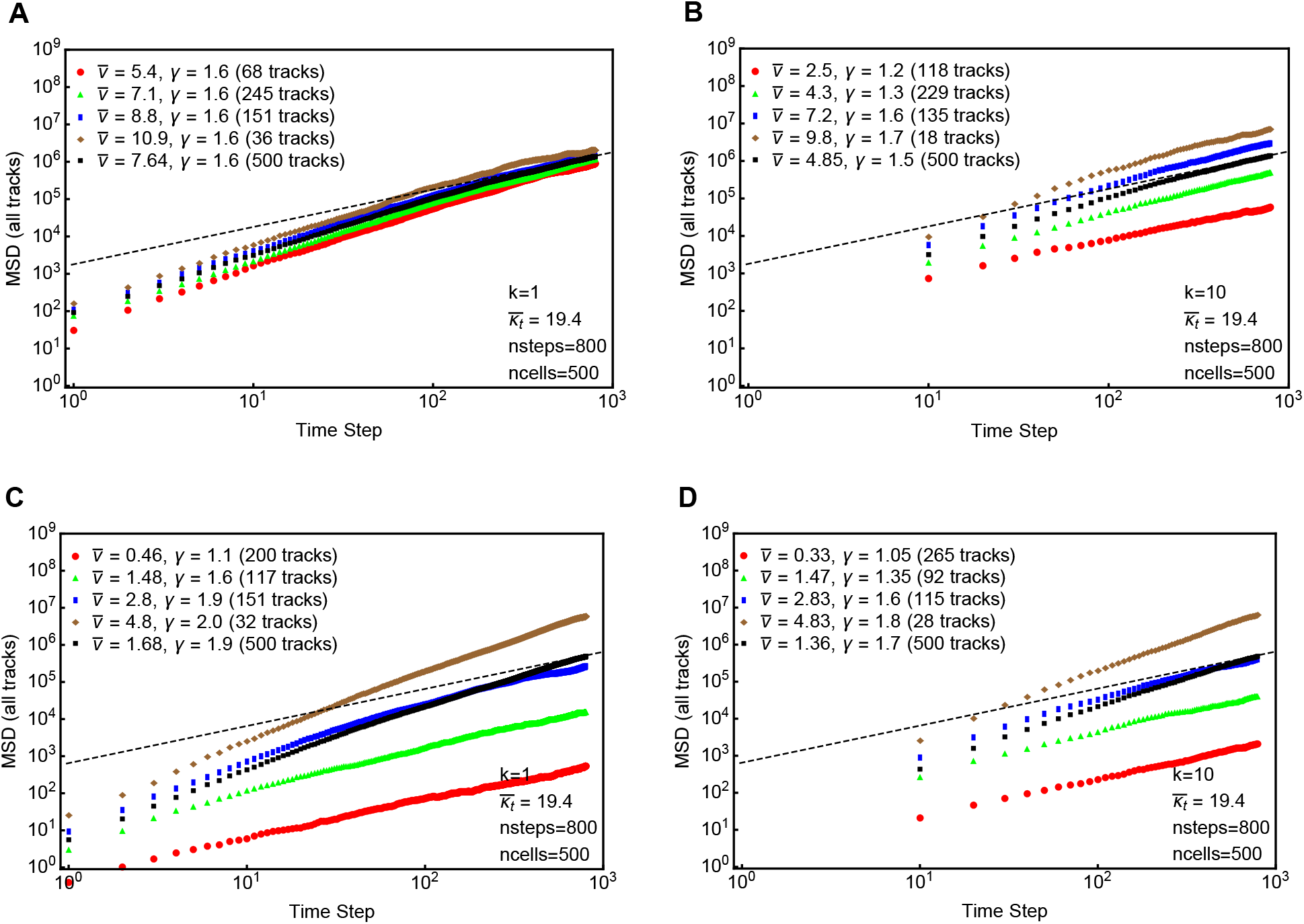
Change in mean square displacement (MSD) with time for different cell cohorts is qualitatively equivalent whether there is or there is not an intrinsic link between the average speed and average turning angle per cell under coarse sampling of the cell trajectories. We simulated movement of cells assuming a correlated random walk characterized by the concentration parameter *κ*_*t*_ of the vMF distribution (see eqn. (1)) and an independent distribution of cell speeds (characterized by the mean 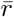 of the Pareto distribution, see eqn. (3); panels A&B) or when there is a direct correlation between *κ*_*t*_ and cell speed (panels C&D). We sampled the movement data either every step (*k* = 1, A&C) or even 10^th^ step (*k* = 10, B&D). In all simulations, the parameter *κ*_*t*_, which dictates the average turning angle of a cell (i.e., persistence), is randomly drawn from a log-normal distribution with mean *µ* = 1 and standard deviation *σ* = 2 (eqn. (4)). The timestep indicates the regular intervals at which we simulated the cell positions, and MSD is dimensionless. In one set of simulations (A-B), we randomly draw 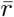 from an independent log-normal distribution with mean *µ* = 2 and standard deviation *σ* = 0.2. In the Pareto distribution we set *α* = 3 for every cell we calculated 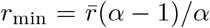. In another set of simulations, we let 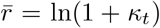) for every cell (C&D). Simulations were done with *n* = 500 cells for 800 timesteps. Four speed bins were considered from the distribution of average speed per cell and for each bin MSD was computed. We also include the MSD for all tracks together. To characterize the MSD we used the relation MSD = *ct*^*γ*^ where *γ* = 1 suggests Brownian diffusion (denoted by a thin black dashed line) and *γ >* 1 suggests superdiffusion. The parameter *γ* was estimated for each MSD curve by linear regression of the log-log transformed MSD data.

In another set of simulations, we let the persistence *κ*_*t*_ be drawn from a lognormal distribution with 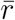 being directly determined by *κ*_*t*_ through the relation 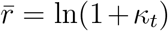. As expected, frequent sampling (every movement, *k* = 1) resulted in cell cohorts with different speeds displaying different rates of displacement, with slower cells displacing nearly as Brownian (*γ* 1, **Figure 5**C). Importantly, for coarsely sampled data (*k* = 10), displacement of cell cohorts with different speeds did not dramatically change (**Figure 5**D), and the curves were similar to the simulations in which speed and persistence were not intrinsically correlated (compare **Figure 5**B&D). This further suggests that MSD plots for cell cohorts with different speeds cannot be used to infer the intrinsic correlation between speed and turning.

### Sub-sampled simulations can match experimental data

So far our simulations did not allow to find a significant difference in several major characteristics of cell movement between a model with an intrinsic link between cell persistence and speed and a model without such link. We therefore wondered if comparing simulations to actual data may allow to see the inadequacy of the “sub-sampling” model. Comparing the model simulations with the data was not trivial, however, because 1) our vMF distribution-based simulations were done using dimensionless units (time step, movement length) and experimental data have dimensions, and 2) specific elements of the data that need to be compared with those of simulations may be debated. We opted for a simple approach whereby we attempted to reproduce the experimentally observed correlation between average turning angle and average speed per cell with that found in simulations. For a fair comparison we needed the data to be sampled at regular time intervals but we found that in about 35% of tracks (out of 712) in control experiments of Jerison & Quake [18] there were missing time step values. We therefore cleaned the data splitting the tracks with missing values and assigning new track ID to each of the new trajectories as we described recently [11]. This resulted in 1337 trajectories with now identically spaced measurements (every 45 sec) with 68 movements per track on average. Importantly, 101 of these had only 1 or 2 time steps that did not allow to calculate the average turning angle and average speed per track, hence these were removed, resulting in 1236 tracks which were analyzed further (**Supplemental Figure S15**). For every trajectory we calculated the average speed and average turning angle and found a strong negative correlation between the two parameters (**Figure 6** and **Supplemental Figure S15**).

**Figure 6:**
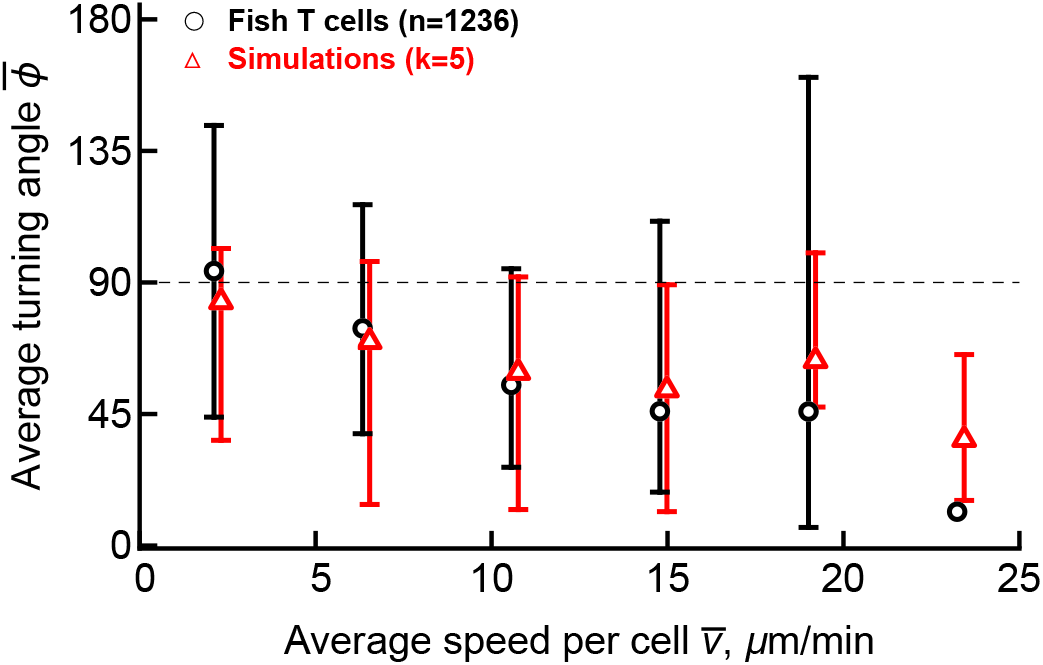
Matching simulations of cell movement in which cell’s speed and turning ability are uncorrelated with experimental data. For cleaned data of Jerison & Quake [18] we calculated the average turning angle and speed for every trajectory and plotted binned data. Note that the bin with the largest average speed had only one trajectory. We also performed simulations in which every cell has a defined persistence ability and speed, sub-sampled the resulting simulation data (every *k* = 5th step was used, see **Supplemental Figure S15** for more detail). To simplify calculations we assumed that this sampling frequency is 1min, calculated the average turning angle and average speed for every trajectory, and then binned these simulation data in the identical way to that of actual experimental data. Confidence intervals denote 2.5 and 97.5 percentiles of the data. Also note that while experimental data were collected in 2D by ignoring z-coordinates of the moving cells [18], our simulations were done in 3D.

We then performed a series of simulations with 1000 cells each undergoing 100 movements by varying the distribution of persistence ability (determined by concentration *κ*_*t*_) and speed (determined by the average movement length 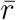) and sub-sampling frequency (*k >* 1). Then for every track in the simulated data we calculated the average speed and average turning angle. Because we could not find solid methods to compare two scattered plots, we binned experimental data and data from simulations into cell cohorts with similar average speeds (**Supplemental Figure S15**). We found that by changing parameters of the distribution for *κ*_*t*_, 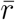 and sub-sampling frequency *k* we could relatively well visually match the experimental data for some parameter combinations (e.g., **Supplemental Figure 6**) further suggesting that a model in which turning ability of T cells and their intrinsic speed are uncorrelated is consistent with experimental data for some parameter values that do not appear to be unrealistic.

The main issue with modeling PRWs with vMF distribution is that all the model parameters are unconstrained by the data. In contrast, Wu *et al*. [16] method of simulating PRWs may be more constrained by experimentally measured parameters such as sampling frequency and persistence time of cells. Because the data on T cell movement in zebrafish are 2D projection of 3D movements (which may generate artifacts), we analyzed another recently published dataset on 3D movement of naive CD8 T cells in murine lymph nodes [11]. For this dataset we found a strong negative correlation between average speed and average turning angle for over 1000 trajectories (**Supplemental Figure S16**A). We could relatively well estimate the persistence time for individual cells in the data (**Supplemental Figure S16**B) with imaging frequency *k* = 20 sec being fixed in the experiment [11, 14]. We randomly assigned every cell in the dataset a speed (**Supplemental Figure S16**C), simulated cell movement using Wu *et al*. [16] method (eqn. (5)) with the time step of 1 sec, and sub-sampled every trajectory with *k* = 20 sec as it was done experimentally. Interestingly, for a specific distribution of speeds given a lognormal distribution we found a strong negative correlation between speed and persistence with magnitude (characterized by Spearman *ρ*) being similar to that observed in the data (**Supplemental Figure S16**A&D). Visually, however, the simulated correlation was not fully matching the experimentally observed correlation with most cells in simulations exhibiting small turning angles and high speeds. This is likely because the chosen lognormal distribution of speeds may not be fully matching to what may be in the data. Our results suggest that while sub-sampling may contribute to the observed correlation between speed and turning, specific elements of that correlation, e.g., mass distribution along the correlation line, may provide ways to falsify the sub-sampling as the main driver of the correlation.

## Discussion

Cell migration is a complicated process. In general, cells move randomly, often by correlated random walks as determined by the turning angle and movement length distributions [11]. Whether there are basic fundamental principles that determine movement strategies of cells remains debated. Recent studies with various cell types, conditions, and constraints (e.g., genetic mutations) have shown that faster cells tend to move straighter (i.e., more persistently) and slower cells tend to change direction more often, resulting in a positive correlation between persistence time and cell speed, or equivalently, in a negative correlation between average turning angle and average speed per cell [18, 26–28, 31]. The generality of this correlation found for different cell types and conditions is a strong argument that correlation arises due to a fundamental, cell-intrinsic movement program that is conserved across different systems [18, 26]. Yet, there are examples of experiments in which speed and persistence are not correlated [25, 32]. For example, Stokes *et al*. [25] found migration of microvessel endothelial cells in control medium with or without agarose overlays resulted in the same persistence times but different speeds.

Here we argue that because turning angles and speeds are measured parameters, they are sensitive to the frequency at which the movement of cells is recorded. The assumption that cells in a given population have a broad and uncorrelated distribution of intrinsic movement speeds and turning abilities naturally results in a negative correlation between average turning angle and average speed when sampling of cell trajectories is done less frequently than the cell’s “decision” for turning (**Figure 2**); such sub-sampling must occur at the frequency that is similar to typical persistence time of cells (**Supplemental Figure S8**). Sub-sampling at frequencies that exceed persistence times of all cells will result in lost correlation between speed and turning (or speed and persistence, **Supplemental Figure S12**). These results are valid both in 3D and in 2D; in the latter case simulations can be done using von Mises distribution to model correlated random walks. We found the same conclusion that sub-sampling of cell trajectories results in negative correlation between speed and turning using an alternative framework to simulate correlated/persistent random walks based on OU process suggesting that our results are not an artifact of vMF-based simulations (**Supplemental Figure S13**). Given that in typical *in vivo* experiments sampling occurs at a frequency that of the same order of magnitude as the typical persistence time (e.g., intravital imaging of T cell movements is typically done at frequency of 1 per min and typical T cell persistence time in LNs or liver is 2-10 min, [11, 20]) sub-sampling is likely to contribute to the observed correlation between speed and turning.

We found that simulating cell movement assuming that there is or is not a link between persistence and speed generated very similar movement characteristics, for example, change in MSD vs. time (**Figure 5**). Furthermore, the average turning angle or average speed were similar in two models when sampling rate was changed (**Figure 4**). Interestingly, we could relatively well match the experimentally observed correlation between average turning angle and average speed per cell using simulations for T cells in fish or in murine LNs (**Figure 6** and **Supplemental Figure S16**). However, matching the data with vMF distribution-based simulations involves several free parameters such as the distributions of turning abilities (determined by *κ*_*t*_) and speeds, and sub-sampling frequency. We found that to match the data we had to assume the actual dimension value for the frequency of sub-sampling. Specifically, to produce the results in **Figure 6** we assumed that sampling occurs every minute (so that movement lengths calculated in simulations are then the cell speed in *µ*m/min). This implies that actual cell movements occur at every 12 sec (*k* = 5), and for such movements, the average movement length is 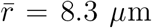. Whether cells are able to make average movements at speeds of 8.3*µ*m/(12*s*) × 60*s/*min = 41.5 *µ*m/min and what is the maximal speed that T cells (or other cells) can exhibit has not been rigorously investigated. In contrast, to match the movement data of naive CD8 T cells in murine LNs using PRW model simulated with the Wu *et al*. method did not require too many assumptions because sampling frequency and persistence time distribution for individual cells were taken directly from the data (**Supplemental Figure S16**).

While it has been typical to infer the correlation between speed and turning by calculating these average characteristics per cell, it is possible to detect changes in speed and persistence for individual cells when tracking is done over extended periods of time, e.g., for amebas [28]. Because Jerison & Quake [18] tracked T cells in zebrafish *in vivo* for extended periods of time we investigated if speed and turning of individual T cells were correlated. For every cell movement we calculated instantaneous speed of the cell and the average turning angle for all the following movements for that cell. We found that out of 1067 trajectories containing 10 or more movements, only 17% had statistically significant correlation between speed and turning angle (after correcting for false discovery rate of 0.05), and 86% of these were negative suggesting limited evidence of correlation between speed and turning for vast majority of T cells *in vivo*.

Our work has several limitations. The major limitation is that we could not come up with an experimental or computational way to reject the null model in which the negative correlation between average turning angle and average speed arises due to sub-sampling of trajectories of heterogeneous cells. This remains a challenge for future studies. Moreover, how sampling (sub- and over-sampling) contributes to the observed patterns of cell movement remains to be investigate more rigorously, and studies that can estimate the “speed-of-light” of cells (i.e., maximal speed that cells can actually have) would be important to constrain impact of sub-sampling on inferred speeds of cells. Studies measuring kinetic details of cell migration suggested that actin, the major protein involved in cell movement, can polymerize at a rate of 5 *µ*m/s = 300 *µ*m/min [33]. Therefore, the potential upper limit of cell’s instantaneous speed is unlikely to exceed this limit.

Another limitation is the relatively arbitrary choice of model parameters, such as distributions of turning angles (characterized by the concentration parameter of the vMF distribution) and distributions of speeds (characterized by the Pareto distribution) and their relationship to the frequency at which movements of simulated cells are sampled. Our results suggest that the negative correlation between turning angle and speed should be observed for some parameter combinations and we found sets of parameters with which we could match experimentally observed correlation (**Supplemental Figure S15**). The assumed distributions did not appear unrealistic. Given the model complexity with 6 parameters (4 parameters for distributions in speed and turning, sampling frequency *k*, and scaling parameter to relate steps to physical units) it is likely that many different datasets can be reasonably well described by the null model but this will require that sampling frequency is at the same order of magnitude as the typical persistence time (**Supplemental Figure S8**). Because our model involves several parameters, it will require a close collaboration with an experimental group to directly determine if it is not possible to find the distribution of parameters and sampling frequency that would not match specific experimental datasets without assuming a direct link between speed and persistence.

Another general limitation that is not limited to our work is the estimation of persistence time of a single cell track. An average turning angle of a cell is a well-defined quantity that can be robustly estimated from the data/simulations. However, estimates of persistence time of individual cells depend on equation fits to the MSD or velocity correlation curves. For single cell tracks the velocity correlation curves can be noisy and highly variable. In particular, we simulated 500 cells with 100 tracks and computed the persistence times for each cell track by fitting an exponentially decaying function to the velocity correlation of each track (**Supplemental Figure S11**). We found a large variation in individual persistence times (see randomly chosen 10 cell tracks in **Supplemental Figure S11**B), however the average velocity correlation curve for all the cells gives a fairly better fit for the persistence time. This wide variation of persistence time estimates are also observed for Fürth equation fits on individual cell MSD curves.

Future studies may need to investigate whether cell-intrinsic correlation between speed and turning impacts predictions on the efficiency of search, for example, of T cells for the infection site (e.g., [29]). One recent study suggested that some level of correlation between speed and persistence length reduces the search time of a target by 10-15% although that result was dependent on specific structural constrains of the environment [34]. Potential metabolic costs of the correlation between speed and persistence and its trade-offs with other cellular processes may need to be included in such calculations of optimality, though.

Previous studies measured how bacterial cells grow in size by tracking the size of individual cells over time with microscopy; by calculating the slope between the relative cell size at cell division and cell division time various models of how cells regulate the time to divide have been proposed (reviewed in [35]). Kar *et al*. [35] challenged these simple regression analyses suggesting that multiple models for cell growth may be consistent with the experimentally measured correlation between cell size and division time; the authors suggested that interpretation of this correlation should be done in terms of a mathematical model of the hypothesized cell division process. Our work suggests a similar approach. For a given specific experimental system where the correlation between speed and turning is observed, authors need to develop alternative mathematical models of the cell movement programs using parameters and imaging setting close to those used in experiments and test if the observed correlation between speed and persistence may arise due to sub-sampling.

Movement persistence, the propensity of moving cells to keep forward movement when environment and conditions are constant, can be postulated as a type of “biological inertia” (or “biological conservation of momentum”) specific to the cells [11]. Per such a postulate, changes in the environment (e.g., shape of the surface on which cells move, cues, or change in nutrients) are responsible for cells changing movement, e.g., to stop and/or turn. It should be noted, however, that such “biological conservation of momentum” or “inertia” should not be confused with physical inertia, because viscous forces are much stronger than the inertial forces for biological (small) cells (the Reynolds number is of the order of 10^*−*4^, which classifies the cell movements under physical processes of low Reynolds numbers [36]). A more integrated experimental approach is needed to be capable of examining continuous cell movements as we have observed in some experiments [15]. By using alternative mathematical models that incorporate different assumptions about cellular motility and sampling close to a specific experimental system, future collaborative studies between experimental and modeling groups should be be able to accurately quantify the impact of sub-sampling to the commonly observed negative correlation between cell speed and turning angle.

## Materials and Methods

### Choosing turning angles using von Mises-Fisher distribution

To simulate cell movement in 3D, we assumed that cells undergo a correlated (persistent) random walk with the degree of persistence determined by the concentration parameter *κ*_*t*_ in the von Mises-Fisher (vMF) distribution [29, 37], which is a probability distribution on an *n*-dimensional sphere (in our case, *n* = 3) that chooses a direction with measurable bias toward a given direction. The distribution is

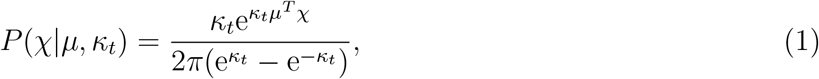

where *µ* is the direction vector toward which there is bias (e.g., the previous movement vector), *χ* is the newly chosen direction vector, *κ*_*t*_ is the concentration (with 0 meaning no bias, positive meaning persistence, and negative meaning aversion), and |*µ*| = |*χ*| = 1. Random (biased) vectors given direction *µ* and concentration parameter *κ*_*t*_ were generated in Mathematica (v. 12.1) by choosing a vector with bias toward direction {0,0,1}, which simplifies the process to choosing 1) *x* and *y* randomly from a normal distribution *N* (0, 1) (using function `RandomVariate`), and 2) *z* based on the von Mises-Fisher distribution, chosen by

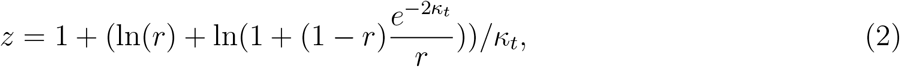

where *r* is chosen uniformly between 0 and 1 [38, 39]. Then *x* and *y* are weighted to place the chosen vector on the unit sphere, and then we use a rotation transform (function RotationTransform) to adjust the generated vector with respect to the desired bias direction. The native Mathematica code to generate a random vector using the vMF distribution is

~~~
vonMisesFisherRandom[\[Mu]_?VectorQ, \[Kappa]_?NumericQ] :=
 Module[{\[Xi] = RandomReal[], w},
 w = 1 + (Log[\[Xi]] + Log[1 + (1 - \[Xi]) Exp[-2 \[Kappa]]/\[Xi]])/\[Kappa];
 RotationTransform[{{0, 0, 1}, Normalize[\[Mu]]}][Append[Sqrt[1 - w^2]
 Normalize[RandomVariate[NormalDistribution[], 2]], w]]]
~~~

### Choosing movement length distribution

The length of the movement *r* was drawn randomly from the Pareto (powerlaw) distribution

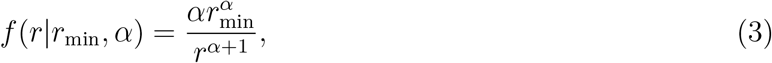

where *r*_min_ and *α* are the scale and shape parameter, respectively, and 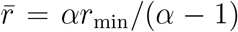. In the Pareto distribution, *r ≥ r*_min_. The Pareto distribution is useful for simulating cell movements because with one parameter, *α*, one can have a thin- or fat-tailed distribution that corresponds to Brownian-like or Levy-like movements [21]. In simulations, we assumed *α* = 5.5, corresponding to a thin tailed, Brownian-like distribution of movement lengths [11], 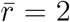, and 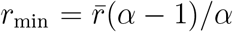. Specific details of the thin-tailed distribution (i.e., distribution with finite mean and variance) are not critical for simulating Brownian-like walks due to central limit theorem [21]. Thus, in our modeling framework *κ*_*t*_ determines the degree of walk persistence (i.e., the average turning angle) and 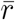 determines the speed of the cell movement. As these quantities are independent, in most of our simulations speed and turning angles truly have no correlation.

### Simulating PRWs using vMF distribution

In simulations, each cell moves in a random direction (determined by *κ*_*t*_ in eqn. (1) and by the previous movement vector) and by a random distance (determined by *r*_min_ and *α* in eqn. (3)). However, if cell movements are measured at a lower frequency than the cell is moving, then the measured cell movement speed and average turning angles are calculated from the “assumed” trajectory that takes into account only some of the cell’s positions. For example, in simulating cell movement for 100 steps where we only count every *k* = 10^*th*^ step as a movement, we calculated instantaneous speeds and turning angles by taking positions 1, 11, … 91 and then calculating the average speed 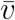 and average turning angle 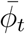 per cell using these positions by dividing the distance travelled by the cell at these time points by *k*. This approach allows to properly compare mean speeds for different sub-sampling rates *k*.

In some simulations, we assumed that every cell has an inherent concentration parameter *κ*_*t*_ which determines cells’ persistence ability. We sampled values of *κ*_*t*_ from a lognormal distribution defined as

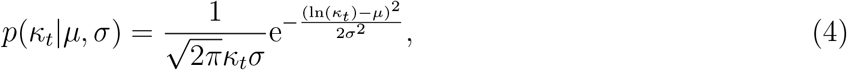

with typically chosen *µ* = 1 and *σ* = 2 to allow for a broad distribution of persistence ability for individual cells (but see Main text for other examples).

We also simulated cell movement with the intrinsic cell movement speed (determined by 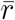) and persistence in the walk (determined by *κ*_*t*_) being correlated. We sampled *κ*_*t*_ for each cell from a lognormal distribution (eqn. (4)) with parameters *µ* = 1 and *σ* = 2 and let the average movement length for each cell be 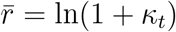. Then, setting *α* = 5.5, we let for every cell 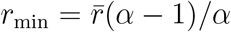 in eqn. (3). Movement length distribution was then sampled randomly from Pareto distribution with *r*_min_ and *α* using `RandomVariate` function in Mathematica.

When simulating a distribution of cell-intrinsic speeds we assumed that for each cell, 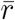 in the Pareto distribution follows a lognormal distribution as for *κ*_*t*_ (eqn. (4)) with *µ* = 1 and *σ* = 2 (and *α* = 5.5). Results were typically independent of these specific parameter values; the only major requirement was that there is sufficient variability in *κ*_*t*_ (or *κ*_*t*_ and 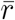) between individual cells.

### Simulating PRWs as OU process

To simulate cell movements in 3D within the framework of OU process we followed the supporting information of Wu *et al*. [16] and Eq. (1) of Jerison & Quake [18]. Movement coordinates of cells at time *t* + *dt* are given by the following equations [16]:

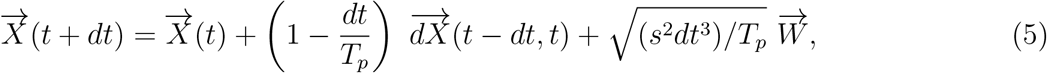

where *T*_*p*_ and *s* are the persistence time and speed, respectively, the position 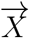 at *t* + *dt* depends on the position at *t* guided by the persistence factor (1 *− dt/T*_*p*_) with a randomness in movement given by the Gaussian noise 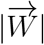 multiplied with a unit magnitude vector, and 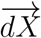 is a displacement vector between times *t dt* and *t*. Given two vectors for subsequent movements, we simulated the next movement vector using a following user-defined function in Mathematica:

~~~
OneMovement[{x1_, x2_}] := {x2, x2 + (1 - dt/pt)*s*Normalize[(x2 - x1)] +
 Sqrt[(s^2*dt^3)/pt]*{RandomVariate[NormalDistribution[0.0,1]],
 RandomVariate[NormalDistribution[0.0,1]],
 RandomVariate[NormalDistribution[0.0,1]]}}
~~~

where `x1` and `x2` are two sequential positions, the displacement with respect to last position is given by the term `(1-dt/pt)`, s is the initial step length to start the simulation, however this s is essentially the speed in *µ*m/min for sampling every step at *k* = 1 sec. Once we have the next movement vector we repeat this for `nsteps` and for `ncells` where `nsteps` is the multiple of `dt=1`sec in our simulations.

### Mean squared displacement

To calculate the mean square displacement (MSD), we used the relation

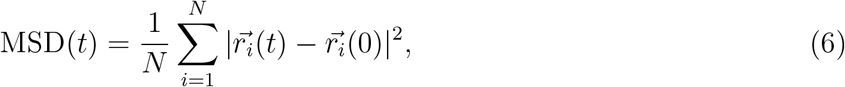

where 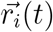 is the position vector of the cell *i* at time *t* and 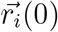 is the initial position and *N* represents the total number of cell tracks considered (typically in our simulations *N* = 500). For different cell cohorts, we divided the total cell tracks into speed groups based on the average speed distribution of the individual cell tracks. To characterize the MSD plots we fitted a relation MSD(*t*) ∝ *t*^*γ*^ using least squares to estimate *γ* for all cells or cohorts of cells with similar movement speeds. Specifically, we log-transformed MSD and time, fitted a line to the transformed data, and estimated the slope *γ*. Note that *γ* = 1 represents normal (Brownian) diffusion and *γ >* 1 represents super-diffusion.

### Persistence time estimation

We have used two different methods for the estimation of persistence time. In the first method we use the Fürth equation

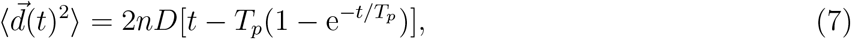

where *D* is the diffusion coefficient of the Ornstein-Uhlenbeck process, *n* is the dimension of the space in which the process is studied and *T*_*p*_ is the persistence time [16, 25]. To avoid potential biases with data selection we fitted eqn. (7) using least squares to all data for the MSD curves.

Second, we also estimated the persistence time by fitting an exponentially decaying function of the form

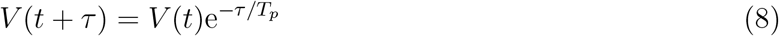

to the velocity correlation curve [24, 25, 30, 40]. Specifically, we calculated the cos *φ* between vectors determining the first movement and every another movement at time step difference *τ* for all cell tracks. We then computed the average cos *φ* at all time steps and plotted them with respect to the time steps. To avoid bias with data selection we fitted eqn. (8) using least squares to all data.

### Persistence time for population average data vs. individual tracks

By the original definition, MSD curves are generated by averaging displacements over cells in the population. However, when enough data is available for individual cells – which is not typical in our experience with *in vivo* imaging data – MSD can be calculated for individual cells [40]. In this approach, squared displacements from the initial position are calculated assuming that “initial” position shifts in time, then the squared displacements can be averaged; however, in this approach average squared displacements at longer time delays are not reliably estimated due to reduced number of data generated. Fürth equation (eqn. (7)) can then be fitted to such data and persistence time for individual cells can be calculated. Velocity correlation curves can be averaged for individual cells in a similar fashion, and persistence time using eqn. (8) can be estimated for individual cells. Note, however, that this approach generates noisy data especially after few time steps that reduces reliability of estimating persistence time for individual cells.

### Parameter dimensions and relating simulations to data

It should be noted that in our simulations with vMF distribution we did not choose dimensions of the parameters, so the time units are in steps (1, 2, 3, etc) and movement length units are also dimensionless. This was done to prevent a bias towards particular numerical values which may differ between different biological systems. Therefore, estimated quantities such as average speed per cell has a dimension of displacement unit per time unit. To link simulation results to experimental data we assumed that when we sub-sample the data at rate *k* (i.e., when *k* = 10 every 10^th^ movement is only used), the calculated average speed per cell is then given in *µ*m/min units. This effectively assumes that if we sample movements every *dt* = 1 min, so the actual movements by cells occur at a rate *dt/k*.

### Statistical analyses

For the correlation between speed and turning (or speed and cell persistence) we performed both Pearson (denoted by *r*) and Spearman rank (denoted by *ρ*) correlation analyses, and in all cases, results were statistically identical. In some cases, the Pearson correlation analysis may not be appropriate due to data not being normally distributed but we used the same test within one set of analyses for consistency.

## Data sources

Data presented in **Supplemental Figure S15** were provided by Jerison & Quake [18] and are available via a link to github in the original publication. Data presented in **Supplemental Figure S16** are from our previous publication [11]; the link to github is provided in the original publication.

## Code sources

All analyses have been primarily performed in Mathematica (ver 12) and codes used to generate most of figures in the paper are provided on GitHub [41]: https://github.com/vganusov/correlated_random_walk_simulations.

## Ethics statement

No animal or human experiments performed.

## Author contributions

The question for the study arose during discussions of the cell movement data between all authors. VVG ran the simulations using vMF distribution-based methods by VSZ and discussed results with other authors. BM ran additional simulations, in particular, based on the PRW model. BM wrote the first draft of the paper. BM, VVG, and VSZ contributed to the final draft of the paper.

## Acknowledgments

We would like to thank E. Jerison and S. Quake for the valuable discussions of our results and comments on the previous versions of the paper that was previously sent as commentary to eLife. This work was supported by the NIH grant (R01 GM118553) to VVG.

## Abbreviations

UCSP: Universal Coupling between Speed and Persistence
MSD: mean squared displacement
vMF: von Mises-Fisher
TA: turning angle
PRW: persistent random walk
OU: Ornstein-Uhlenbeck
LNs: lymph nodes

## Supplemental Information

**Supplemental Figure S1:**
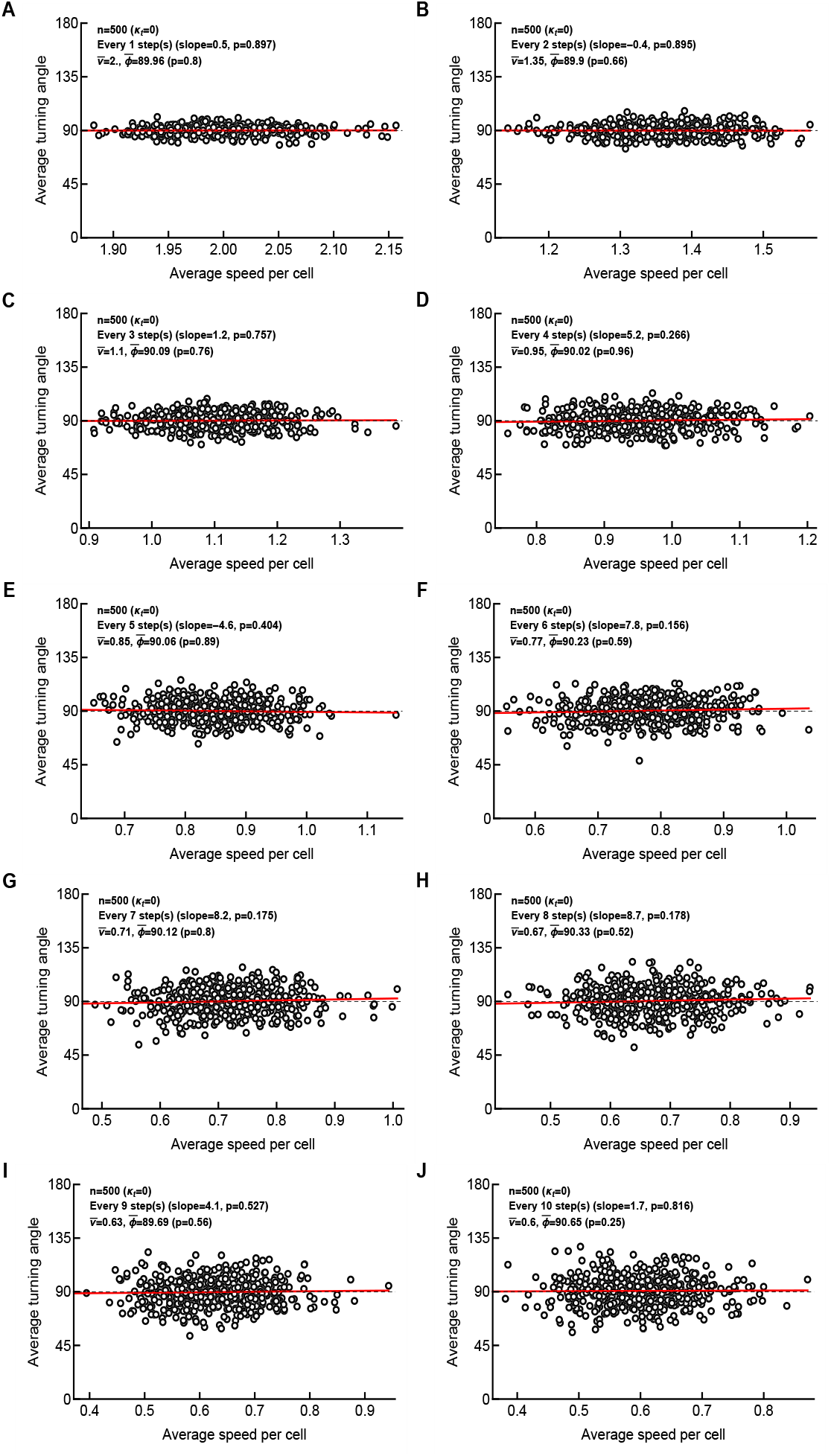
A correlation between average turning angle and average speed does not arise for coarsely sampled data for uncorrelated random (Brownian) walk. Here all cells have the same concentration parameter *κ*_*t*_ → 0 (see eqn. (1)). We simulated movements of 500 cells for 100 steps and calculated the average speed and average turning angle per cell when sampling the data at different frequencies, starting with every step (panel A) and finishing with every 10 steps (panel J). Each panel contains information on the average speed for all cells 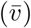, average turning angle for all cells 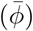, and the result of linear regression of the average speed per cell and average speed per cell (denoted as “slope”) with *p* value from the t-test. We also provide a test if the average turning angle of cells in the population is different from 90^*°*^ (Mann-Whitney test).

**Supplemental Figure S2:**
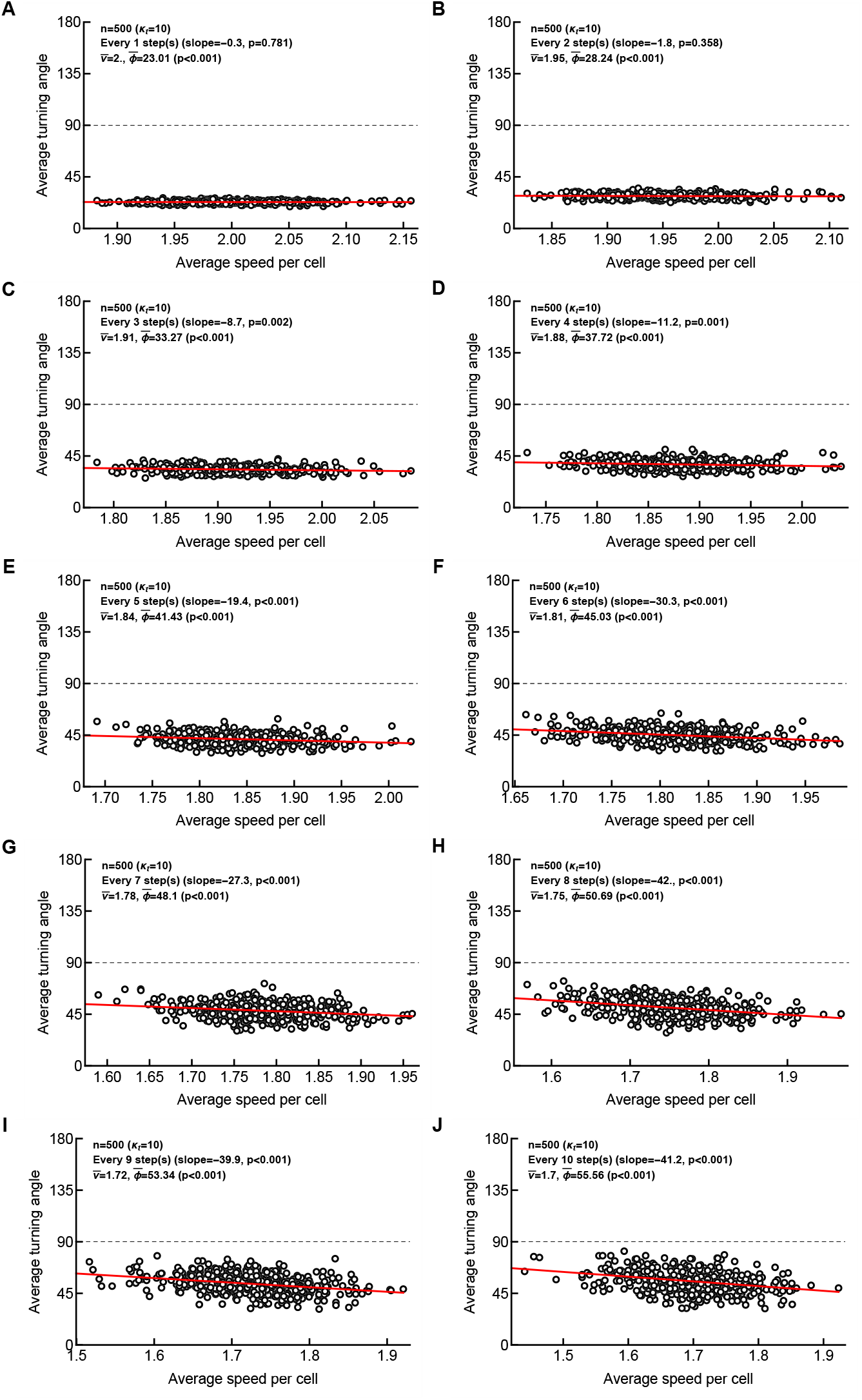
A correlation between average turning angle and average speed appears for coarsely sampled data when cells undergo a correlated random walk. Here all cells have the same concentration parameter *κ*_*t*_ = 10 - note different scale on *x* axis in different panels. See **Supplemental Figure S1** for other details.

**Supplemental Figure S3:**
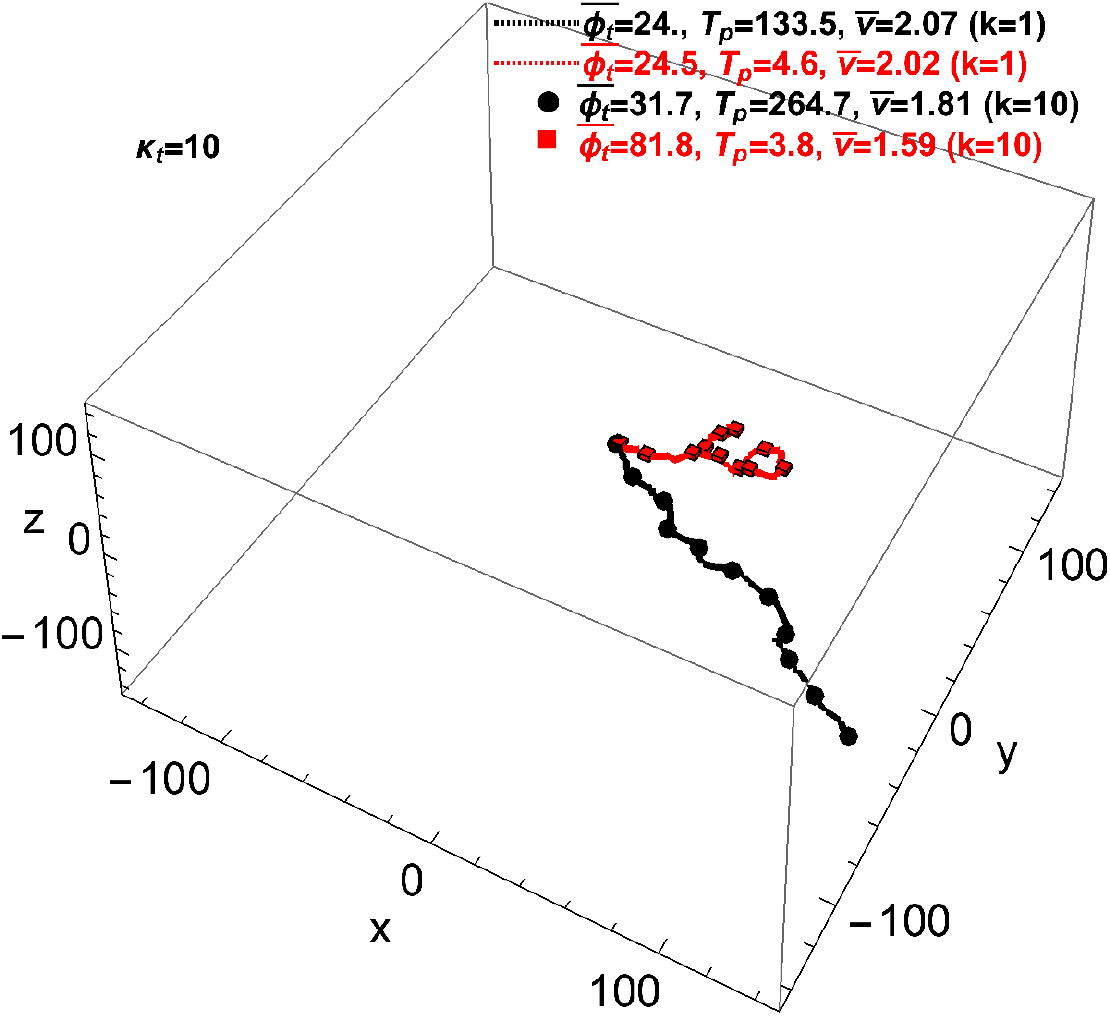
Example of two simulated cells with different trajectories. From simulations in **Supplemental Figure S2** with a fixed *κ*_*t*_ = 10 we sampled two cells with different behaviors. We estimated the average turning angle 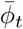, persistence time *T*_*p*_ (from velocity correlations), and average speed 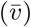 for the data sampled every step (*k* = 1, given by lines) or every 10th step (*k* = 10, given by markers).

**Supplemental Figure S4:**
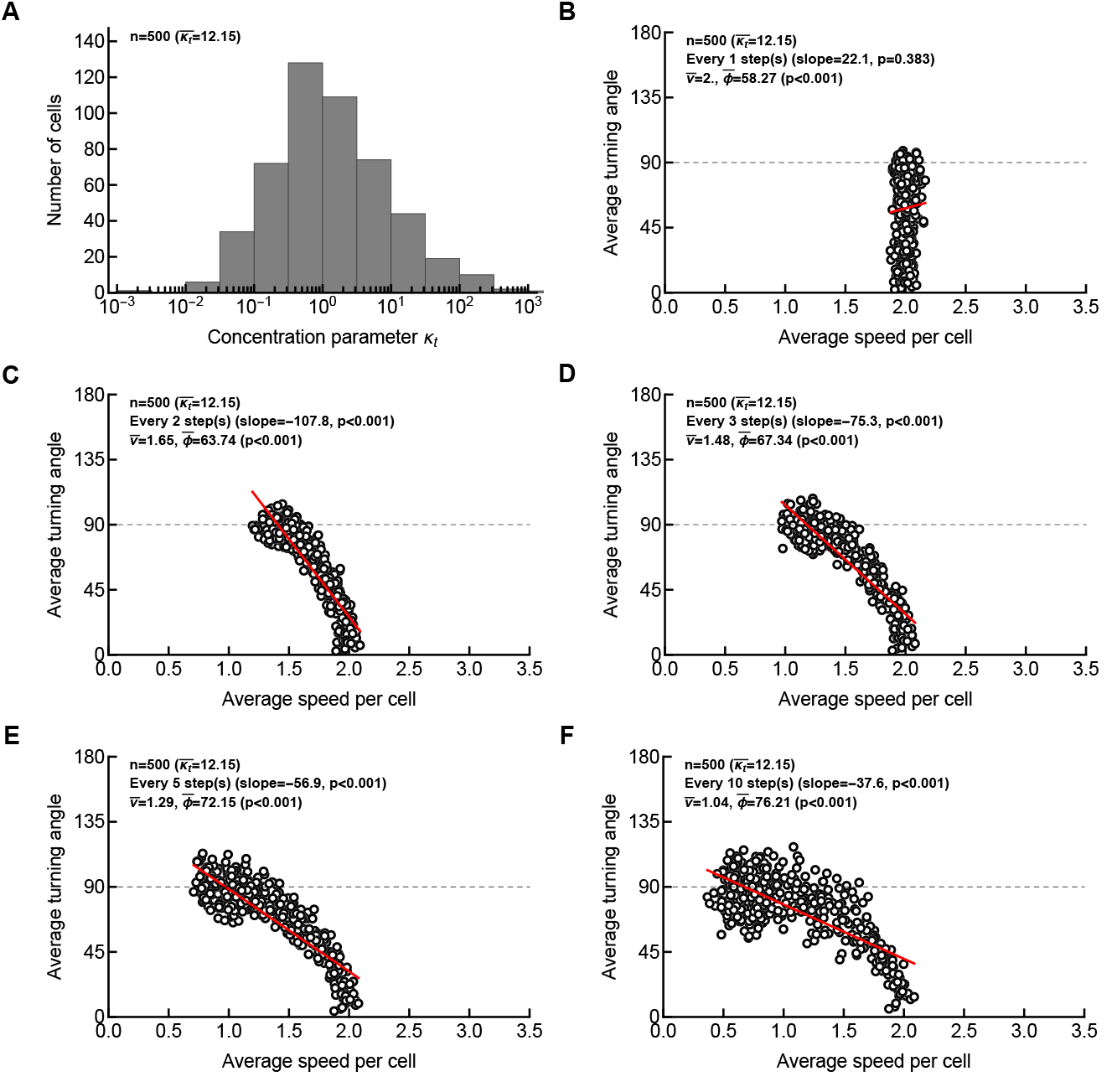
A strong negative correlation between average turning angle and average speed naturally arises for heterogeneous cell populations with coarsely sampled data. We assume that every cell in the population has a different *κ*_*t*_ (from vMF distribution, see eqn. (1)) which was drawn from a lognormal distribution (eqn. (4) with *µ* = 1 and *σ* = 2, shown in panel A), and the movement data were sampled at different frequency (*k* = 1, 2, 3, 5, 10 for panels B-F, respectively). The average concentration parameter 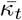 is shown on other panels B-F. See **Supplemental Figure S1** for other details.

**Supplemental Figure S5:**
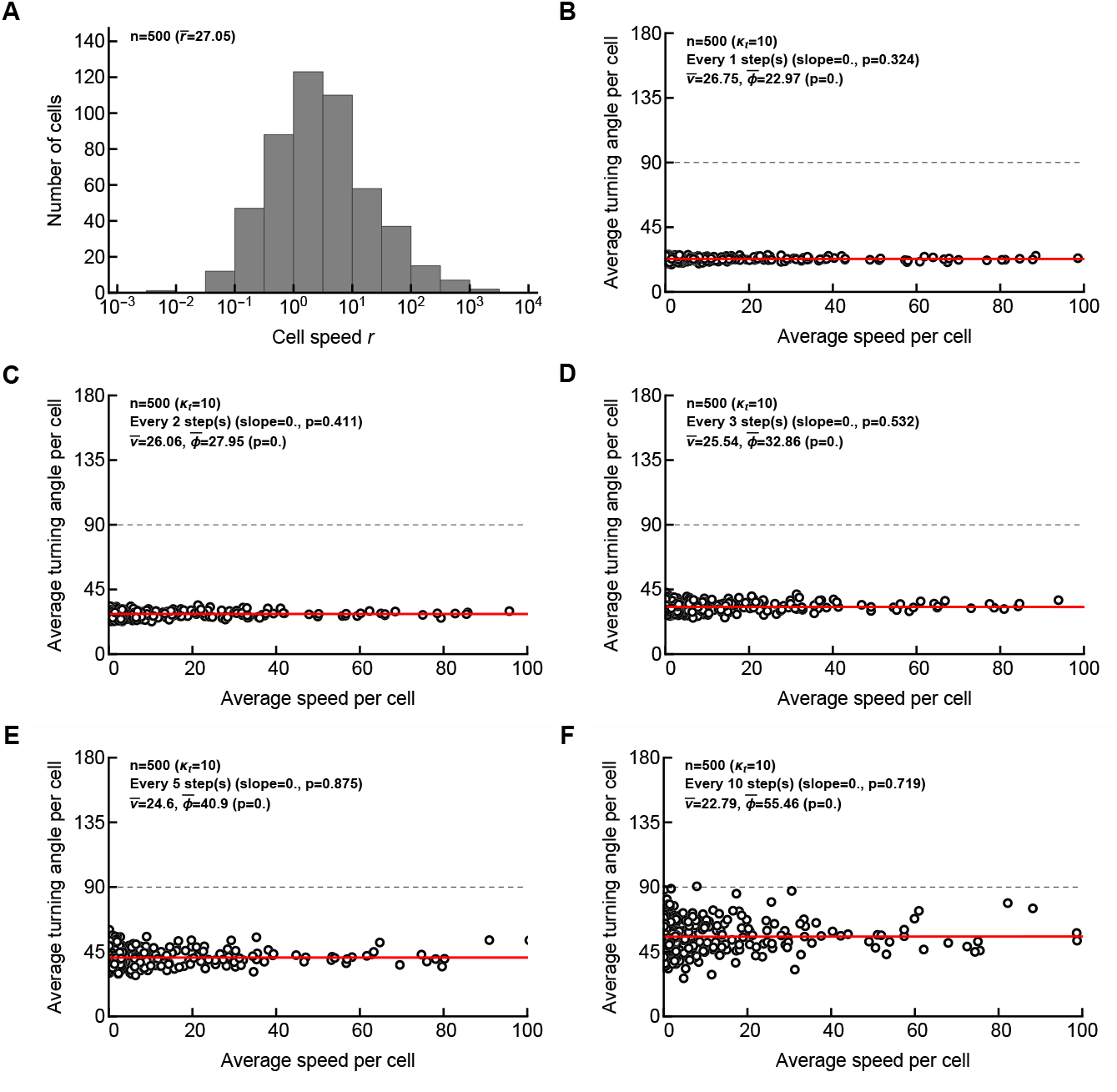
Variability in intrinsic movement speed does not result in a significant correlation between average turning angle and average speed. Here we assume that every cell in the population has a different 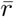 which was drawn from a lognormal distribution (eqn. (4) with *µ* = 1 and *σ* = 2, shown in panel A), and the movement data were sampled at different frequency (*k* = 1, 2, 3, 5, 10 for panels B-F, respectively). Turning angles are defined with *κ*_*t*_ = 10 in vMF distribution (eqn. (1)). See **Supplemental Figure S1** for other details.

**Supplemental Figure S6:**
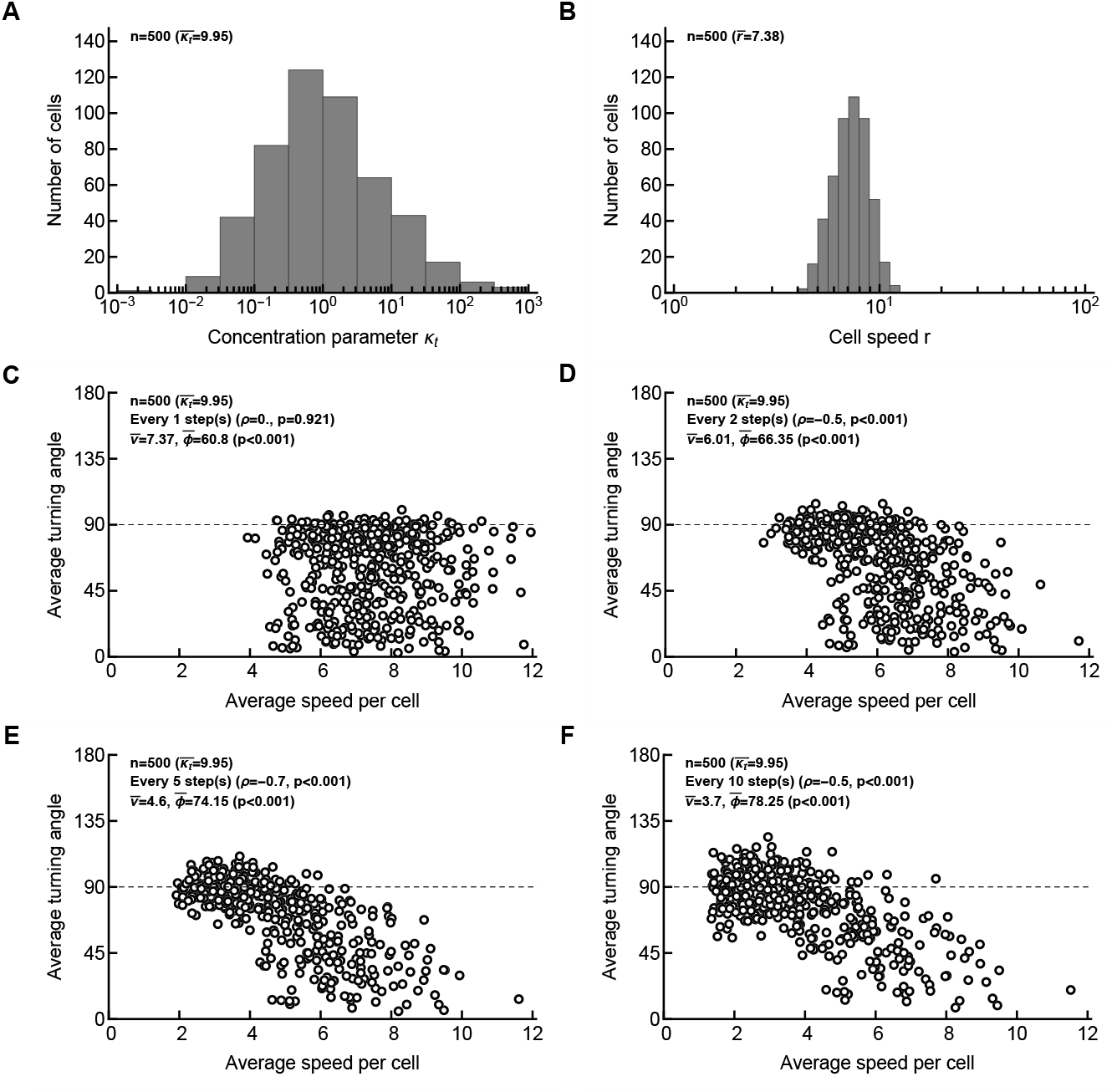
A strong negative correlation between average turning angle and average speed naturally arises when both persistence and cell speeds are variable (but uncorrelated) for different cells for coarsely sampled data. Here we assume that every cell in the population has a different *κ*_*t*_ which was drawn from a lognormal distribution (eqn. (4) with *µ* = 0 and *σ* = 2, panel A), and every cell has a random speed determined by 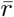 in the Pareto distribution (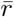 was drawn from a lognormal distribution with *µ* = 2 and *σ* = 0.2, panel B), and the movement data were sampled at different frequency (*k* = 1, 2, 5, 10 for panels C-F, respectively). Correlation was accessed using Spearman rank test with *ρ* and p values indicated on individual panels (because Pearson correlation was not appropriate in this case due to non-normality of the data). See **Supplemental Figure S1** for other details.

**Supplemental Figure S7:**
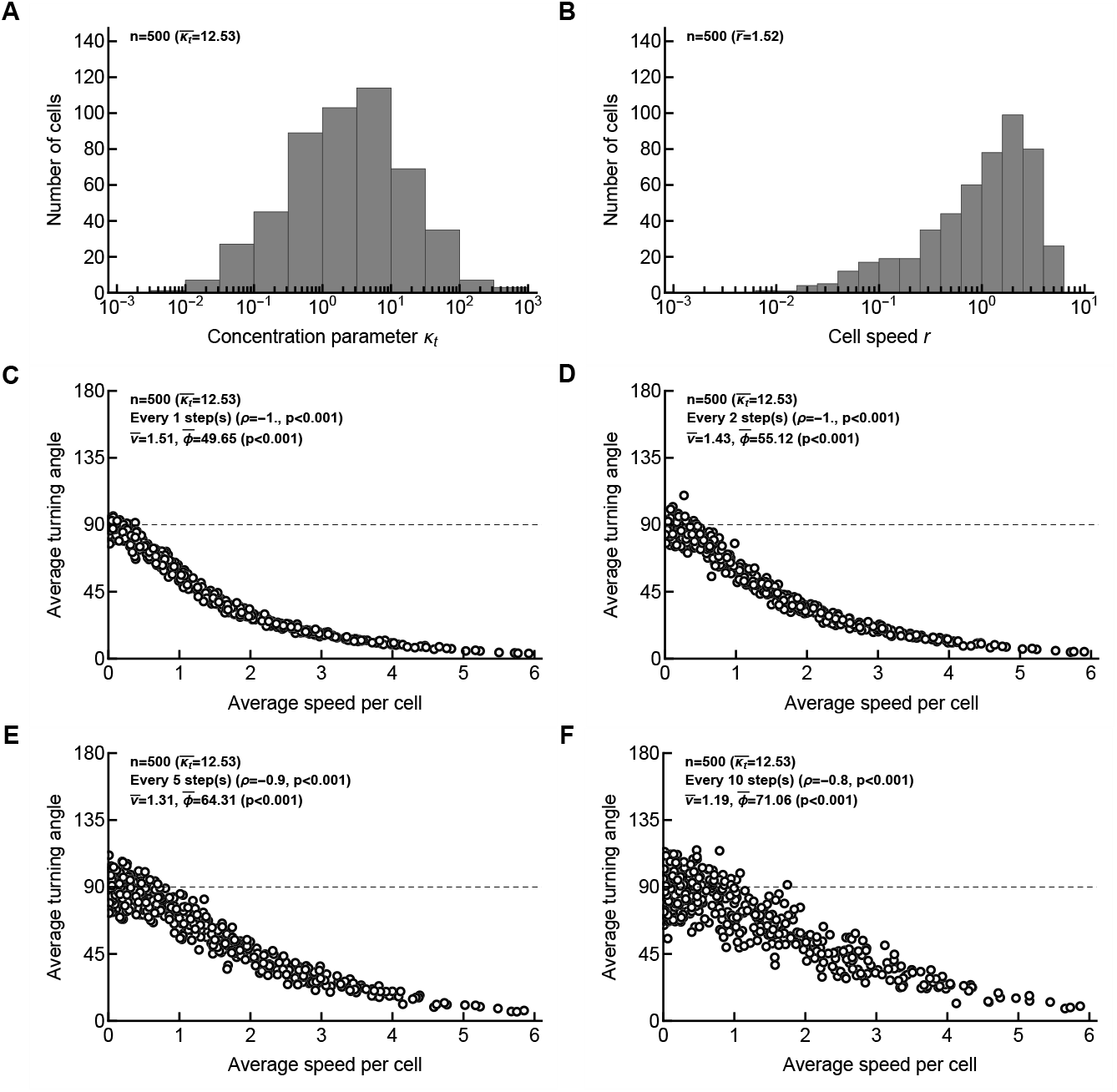
Intrinsic correlation between instantaneous cell speed and turning angles is relatively insensitive to sampling frequency. Here we assume that every cell in the population has a different *κ*_*t*_ which was drawn from a lognormal distribution (eqn. (4) with *µ* = 1 and *σ* = 2, panel A), and every cell has a speed determined by *κ*_*t*_ (i.e., assuming a relationship for each cell as 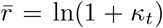 for the Pareto distribution (eqn. (3)), panel B), and the movement data were sampled at different frequency (*k* = 1, 2, 5, 10 for panels C-F, respectively). Correlation was accessed using Spearman rank test with *ρ* and p values indicated on individual panels. See **Supplemental Figure S1** for other details.

**Supplemental Figure S8:**
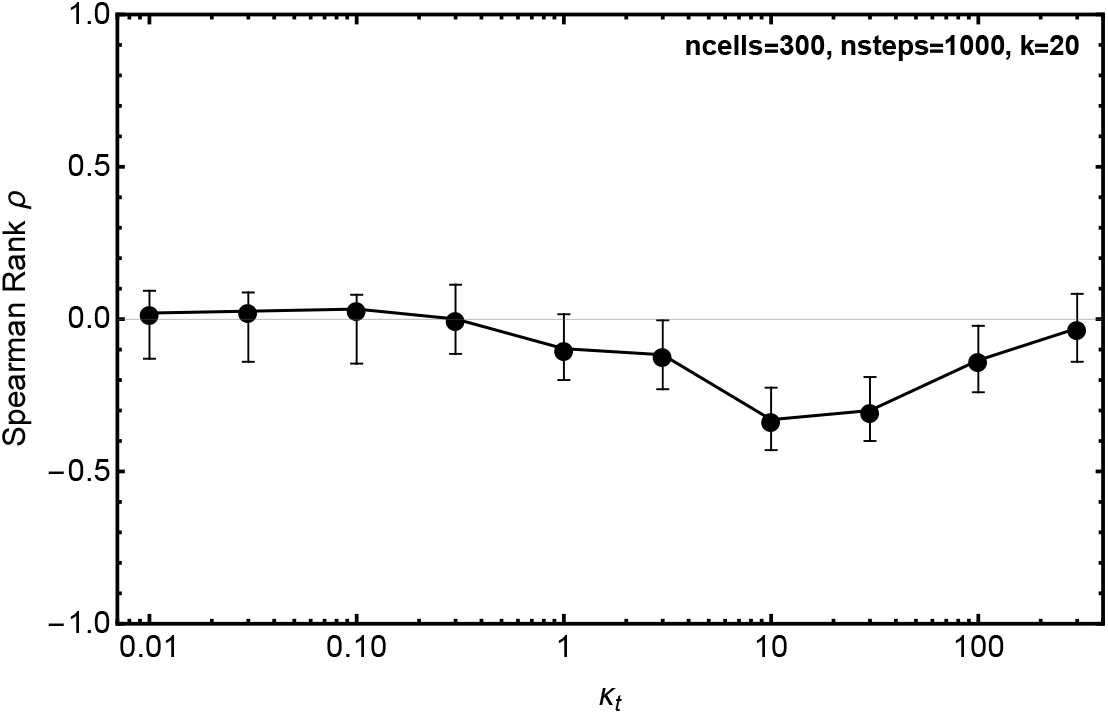
Negative correlation between speed and turning angle arises for a relatively large range of movement persistence (defined as *κ*_*t*_ in vMF distribution-based simulations). We simulated movement of 300 cells for 1000 steps using vMF distribution assuming the same *κ*_*t*_ and speed 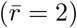 for every cell. We sub-sampled the cell trajectories with *k* = 20. Confidence intervals (95%) for the estimated Spearman rank correlation coefficient *ρ* were calculated using bootstrap approach in routine `SpearmanRho` in `R` package `DescTools`.

**Supplemental Figure S9:**
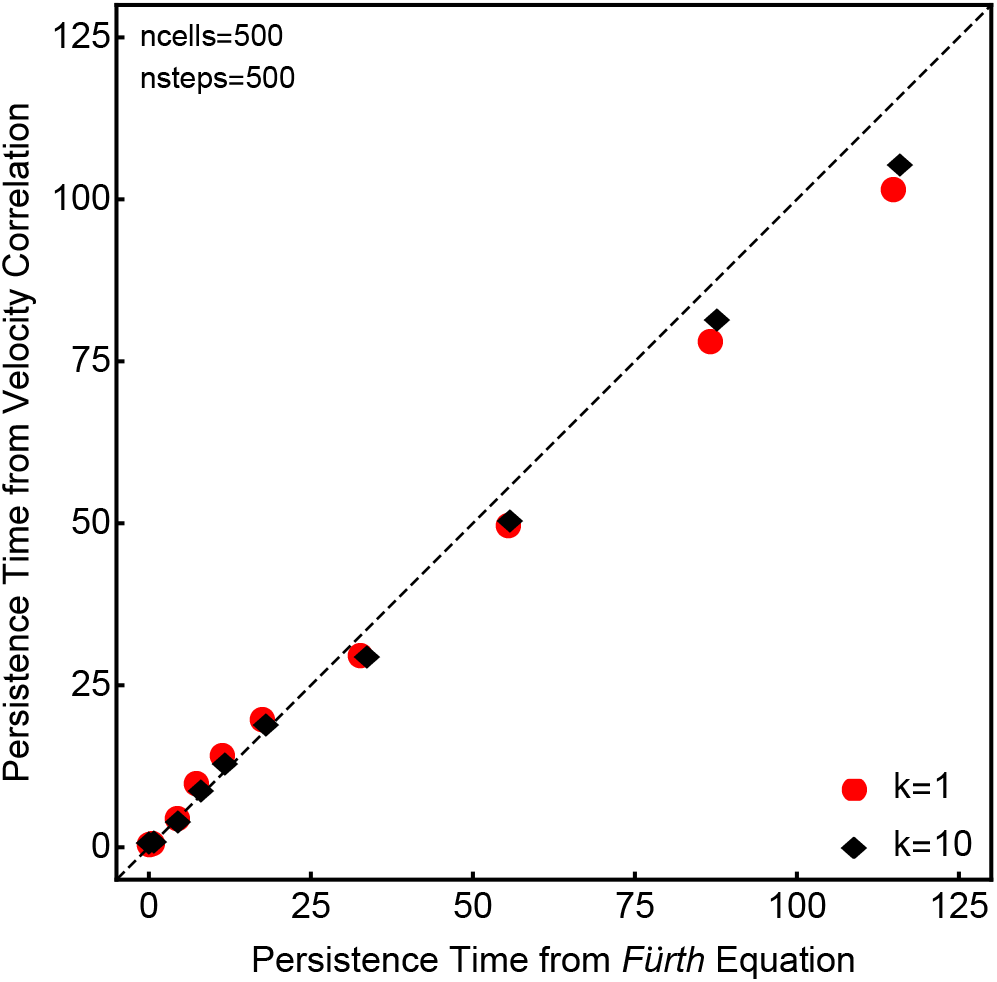
Using Fürth equation to MSD data and fitting an exponential decay function to velocity correlations result in similar persistence times in simulations. We simulated sets of 500 cells each with 500 movements with fixed 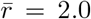 displacement per movement, varied the concentration parameter *κ*_*t*_ in the range 0.1 − 100 for each simulation, and sampled the resulting simulation data every movement (*k* = 1) or sub-sampled every *k* = 10^th^ movement. We estimated persistence time for the whole cell population by fitting the Fürth equation (eqn. (7)) to the population average MSD data or by fitting an exponential equation (eqn. (8)) to the velocity correlation curve of all cells (see Materials and Methods for more detail).

**Supplemental Figure S10:**
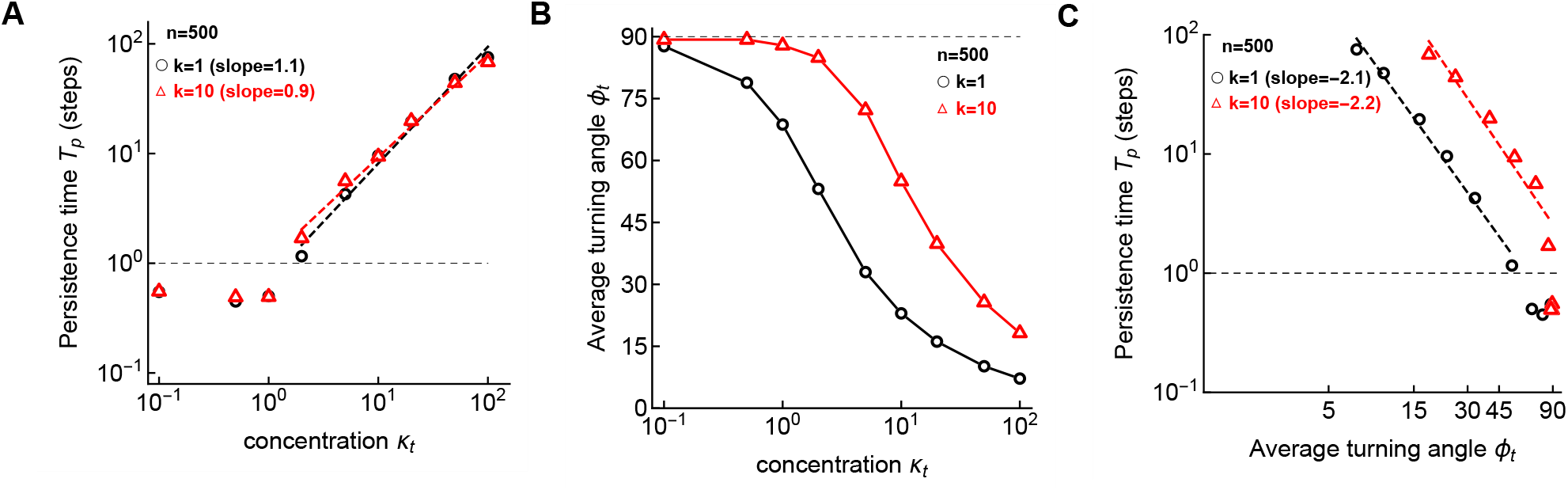
Negative correlation between estimated persistence time and average turning angle. We simulated 9 sets of 500 cell movements each with 100 steps. For the simulated sets we fixed 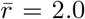 but varied the concentration parameter *κ*_*t*_ of the vMF distribution in the range 0.1 − 100. For sampling the data at every step (*k* = 1) or every *k* = 10^th^ step we then estimated the persistence time (*T*_*p*_) by fitting the Fürth equation (eqn. (7)) to the MSD curves calculated using population average (see Materials and Methods for the estimation of persistence time). We also computed the average tuning angles (*φ*_*t*_) for each set of simulations. In panel A we show the variation of estimated persistence time with concentration parameter *κ*_*t*_, in panel B we show the variation of average turning angle with concentration parameter *κ*_*t*_, and in panel C we show the variation of estimated persistence time with average turning angle. Dashed horizontal line denotes the limit of detection of the persistence time and the data below detection limit were excluded from the regression analysis (shown by the dashed lines).

**Supplemental Figure S11:**
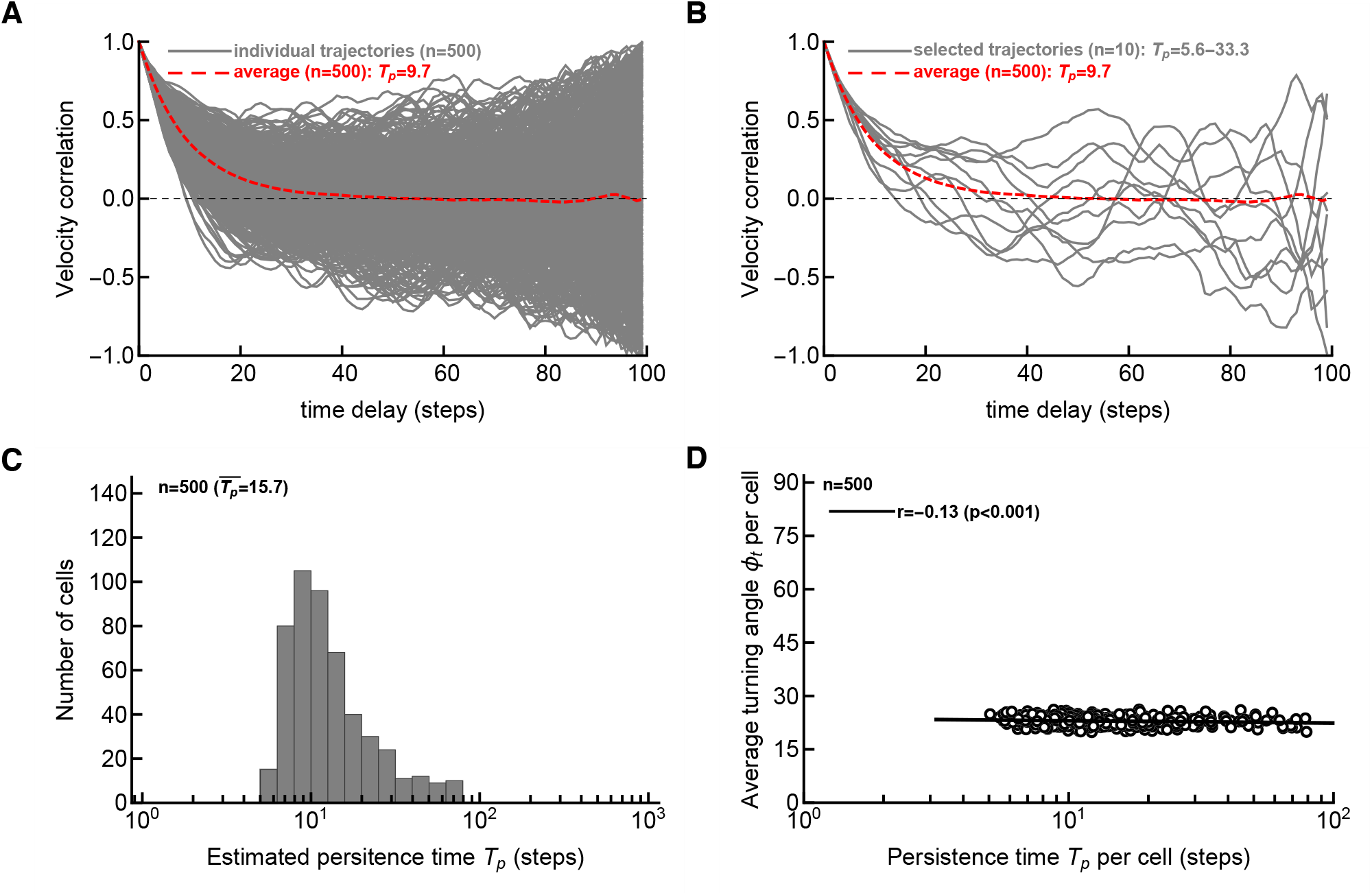
Variability in estimated persistence time for individual cells using velocity correlation. We simulated 500 cells with 100 steps with *κ*_*t*_ = 10 and 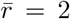 and calculated the velocity correlation curves for data sampled every movement (*k* = 1). By fitting an exponential decay equation (eqn. (8)) to the trajectory data for individual cells we estimated the persistence time *T*_*p*_ for each cell. The red line is the average cos *φ* for all the tracks. In panel B we randomly selected 10 tracks and indicate estimated *T*_*p*_ for these tracks. In panel C we show the distribution of persistence times *T*_*p*_ for all trajectories. In panel D we show statistically significant but weak correlation between average turning angle and persistence time per cell. Note that in these simulations all cells have identical assumed persistence (defined by *κ*_*t*_).

**Supplemental Figure S12:**
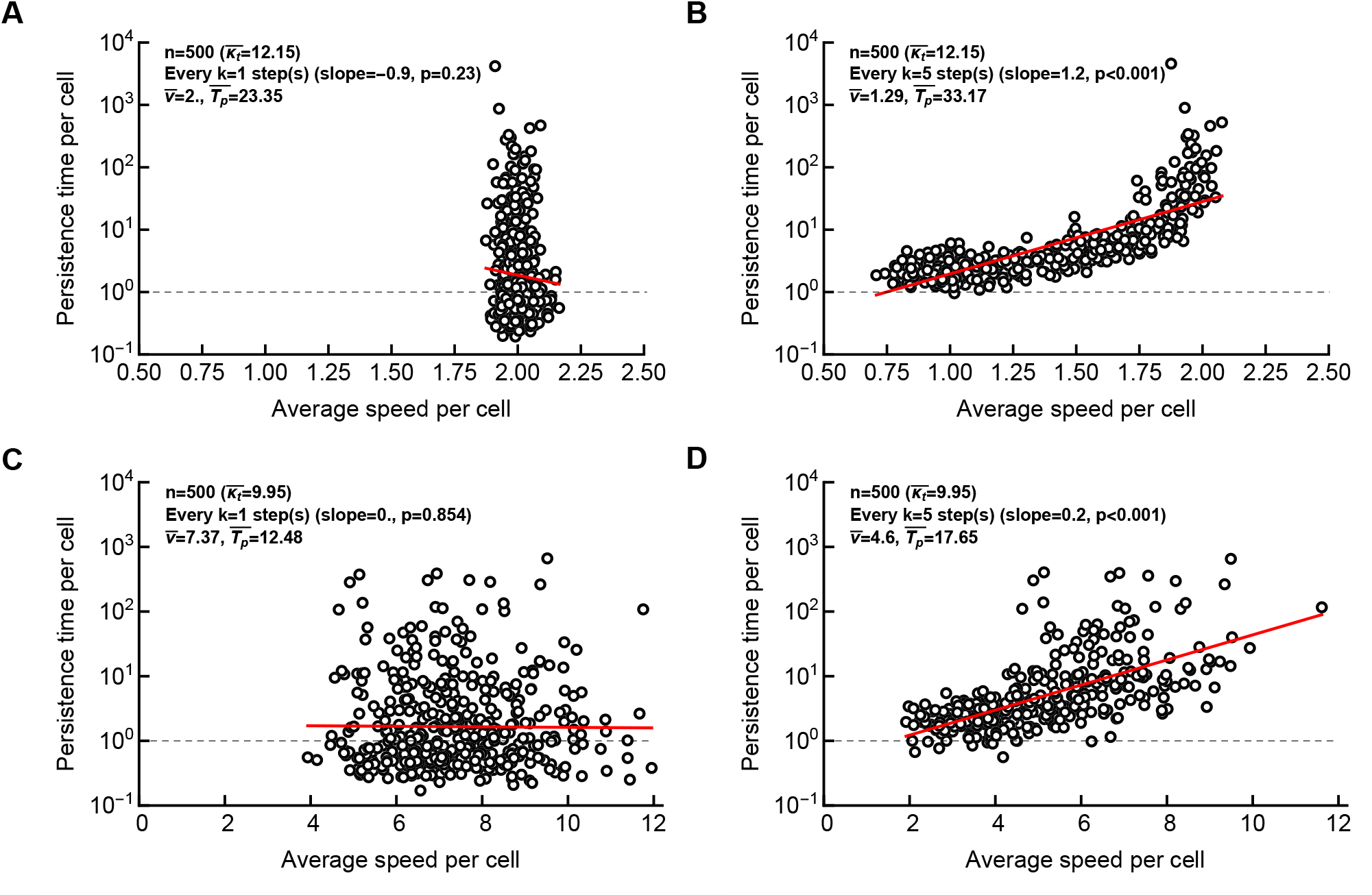
Correlation between persistence time per cell and average speed per cell arises due to sub-sampling. We simulated movements of cells as described in **Figure 2**E-F with *κ*_*t*_ chosen from a lognormal distribution and 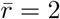 was fixed (A-B) or when 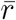 was also chosen from lognormal distribution (C-D); parameters were the same as in **Figure 2**E-F. For each cell we estimated the average speed per cell and the persistence time by fitting eqn. (8) to the velocity correlation curve (see Materials and Methods for more details). In panels A&C sampling is done of every step (*k* = 1) and in panels B&D sampling is done at every *k* = 5^th^ step. Of note, at *k* = 10, correlation between persistence time and speed remained statistically significant, however, we estimated low (*<* 1 step) persistence time for a small fraction of cells which is unrealistic.

**Supplemental Figure S13:**
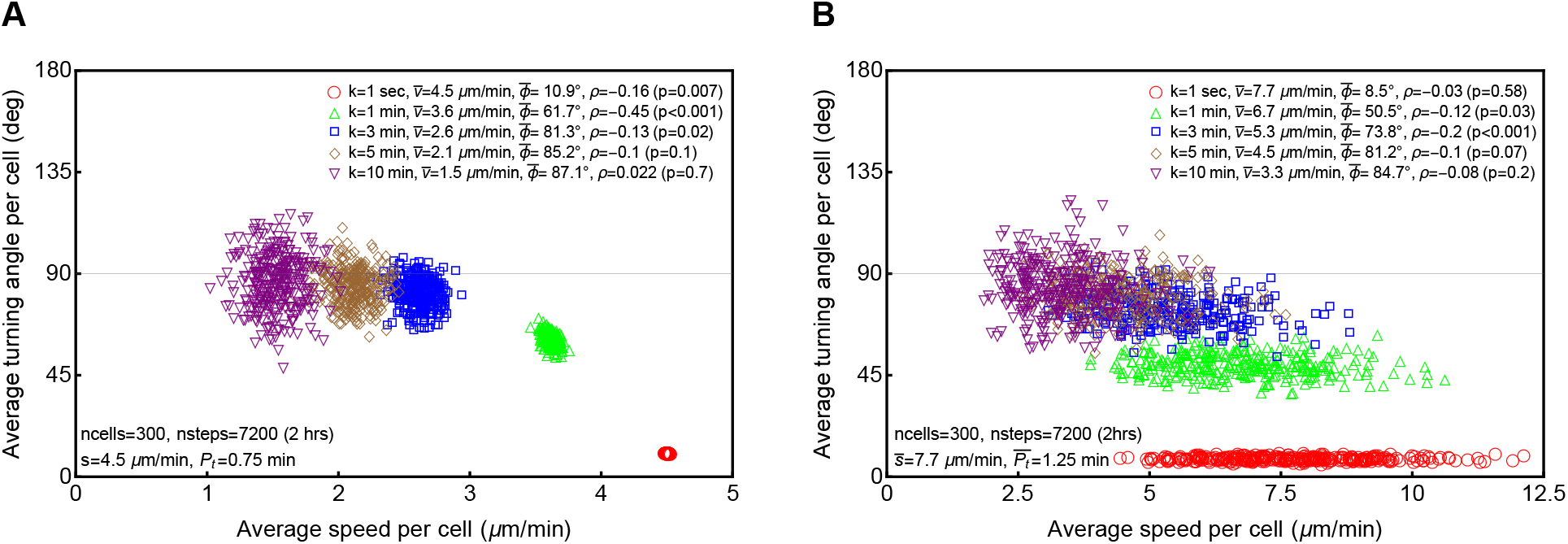
In PRWs simulated as a OU process correlation between speed and turning angle vanishes when sampling interval is much larger than the typical persistence time. We simulate the persistent random walk using a model based on OU framework and described by Wu *et al*. [16]. In panel A we assume that every cell have the same persistence time *T*_*P*_ = *P* = 0.75 min and same speed *s* = 4.5 *µ*m/min. We simulated 7200 steps equivalent to 2 hours for 300 cells and sampled for every *k*^*th*^ min as shown in the legends (1, 10, 50, 100, 200). In panel B we assume that every cell in the population has different persistence times which was drawn from a lognormal distribution (eqn. (4) with *µ* = 0.2 and *σ* = 0.2), and every cell has a random speed drawn from a lognormal distribution with *µ* = 2 and *σ* = 0.2. We simulated 7200 steps equivalent to 2 hours for 300 cells and sampled for every *k*^*th*^ min as shown in the legends. For every set of simulations we also show the average speed 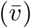, average turning angle 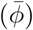 per population, and Spearman rank correlation coefficient (*ρ*) and p value from the test (that *ρ* = 0).

**Supplemental Figure S14:**
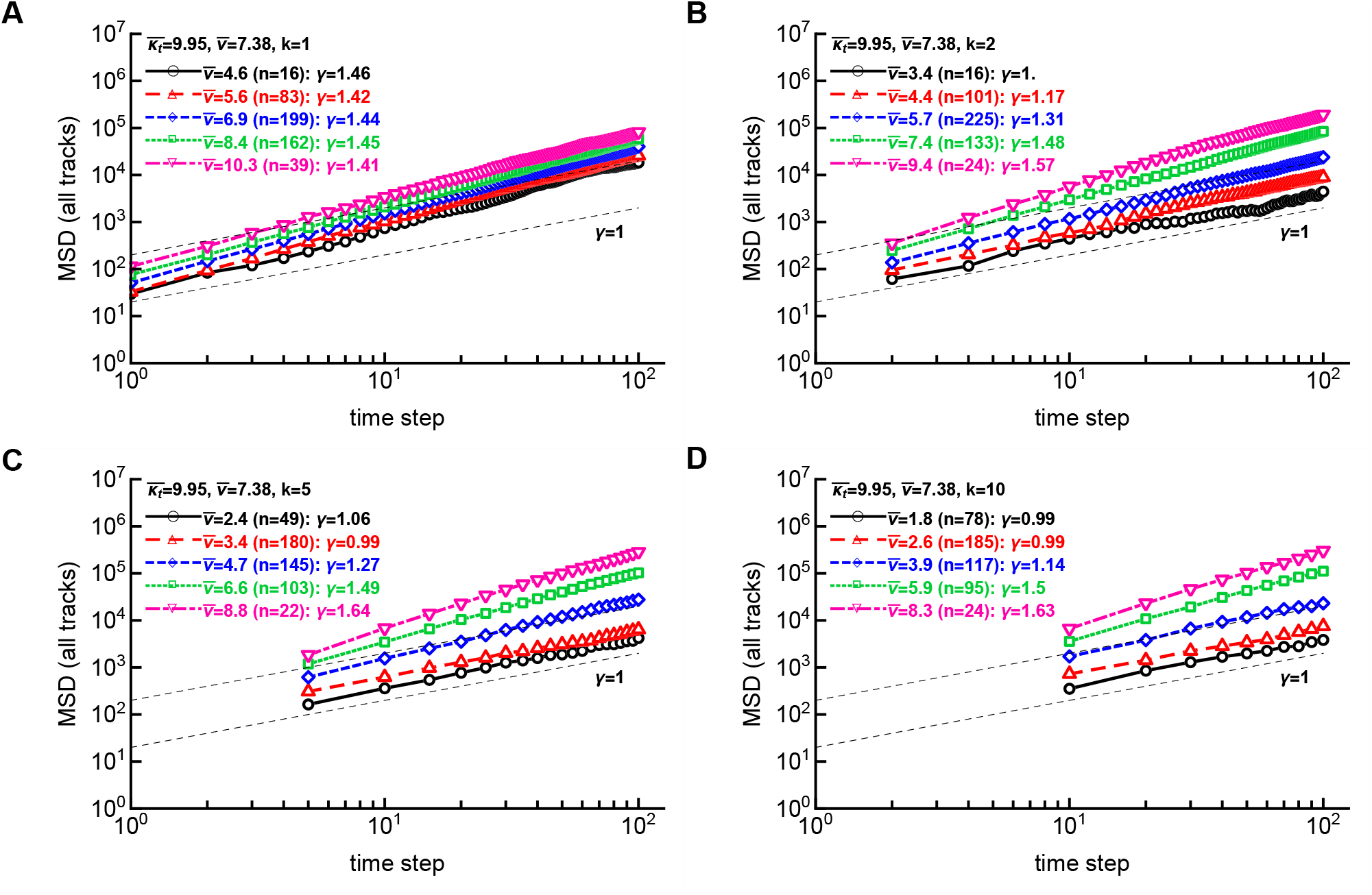
Coarse sampling results in different mean square displacements of cell cohorts with similar inferred average speeds. Here we assume that every cell in the population has a different *κ*_*t*_ which was drawn from a lognormal distribution (eqn. (4) with *µ* = 0 and *σ* = 2), and every cell has a random speed determined by 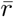 in the Pareto distribution (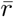 was drawn from a lognormal distribution with *µ* = 2 and *σ* = 0.2). This was identical to how simulations were done in **Figure S6**. We binned the resulting cell trajectories by calculating average speed per track (given specific sampling frequency given by *k*, shown in individual panels) and binned tracks into cohorts with different average speeds. Binning was done by log-transforming the average speeds and then selecting tracks using equally spaced boundaries with 5 bins between minimal and maximal average speeds recorded per sampling simulation. The resulting number of cell trajectories per bin is denoted by *n* in individual panels, together with the average speed of cells in a bin (*v*), and the slope *γ* at which log MSD is changing with log time. Simulations were done with 500 cells for 100 timesteps. Thin dashed lines have a slope *γ* = 1.

**Supplemental Figure S15:**
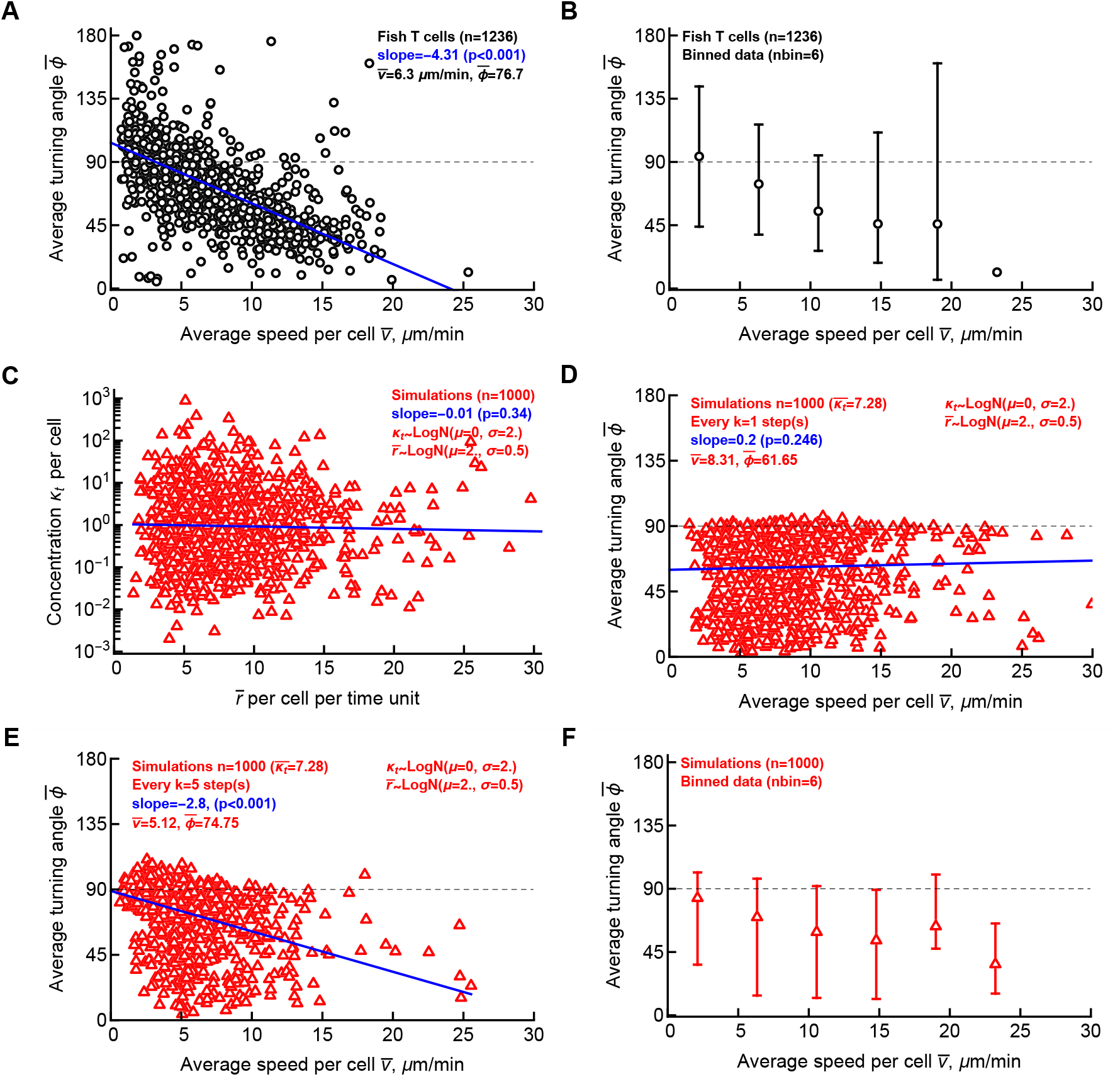
Simulations of cell movement are consistent with experimental data when in simulations cell’s speed and turning ability are uncorrelated but when trajectories are sub-sampled. We cleaned the original data for movement of T cells in zebrafish [18] by splitting cell tracks that had missing coordinates so that every coordinate measurements occurred in equally spaced intervals (45 sec, see Materials and Methods for more detail). For *n* = 712 original cell tracks this resulted in 1236 tracks. For every trajectory we calculated the average speed and average turning angle (A) or binned the data into 6 cohorts with following bin boundaries (0., 4.22, 8.45, 12.67, 16.90, 21.12, 25.34) *µ*m/min (B). We then performed stochastic simulations of 1000 cells for 100 time units with each cell having a defined persistence ability (defined by *κ*_*t*_) and speed defined by 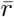, each drawn from a lognormal distribution (eqn. (4)) with parameters *µ* = 0, *σ* = 2 for *κ*_*t*_ and *µ* = 2, *σ* = 0.5 for 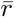, respectively (C). For every track we then calculated the average speed and average turning angle when the data were sampled every time step (*k* = 1, D) or every *k* = 5 steps (E) assuming that sampling in both cases occurs with frequency of 1min. For the simulations data in E we also binned the data the same way as for experimental data (F). Confidence intervals in B and F denote 2.5 and 97.5 percentiles of the data. Other characteristics shown on individual panels are for average speed 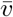, average turning angle 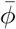. P values for the regression slopes (denoted by solid blue lines) were calculated using linear regression. Another set of parameters for which we found a good match between simulations and data are for *κ*_*t*_ with *µ* = 0 and *σ* = 2 and for 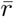 with *µ* = 2.7 and *σ* = 0.2 sampled at *k* = 5 (results not shown).

**Supplemental Figure S16:**
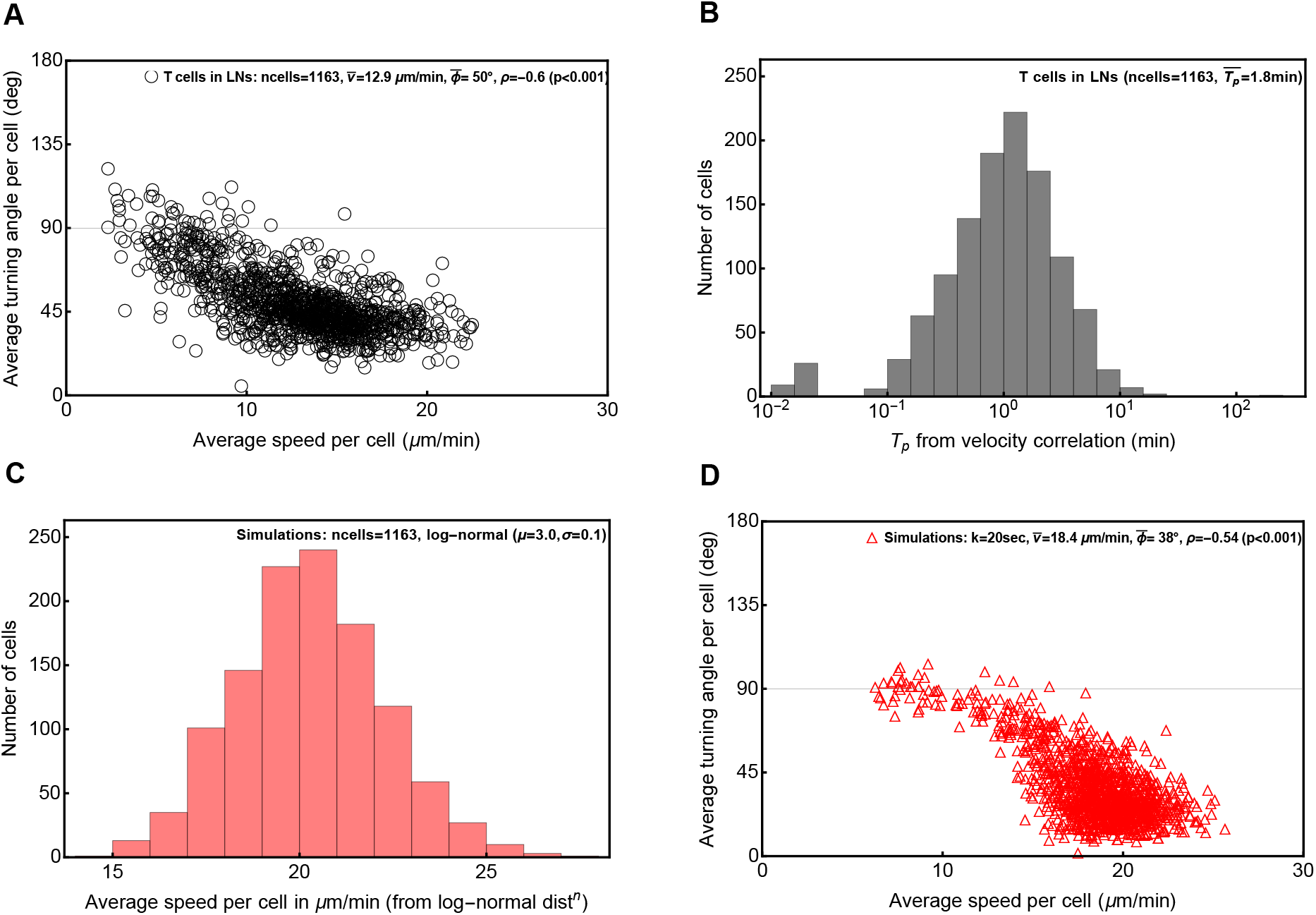
Sub-sampling of simulated cell movements can match experimental data on T cell movement in murine lymph nodes (LNs). For each trajectory in the experimental data on movement of naive CD8 T cells in LNs [11, 14] we calculated the average speed and average turning angle (panel A) or the persistence time from velocity correlations (panel B). We then simulated movement of cells using Wu *et al*. [16] method by taking specific values of persistence time *T*_*p*_ for each cell in panel B and randomly assigning the speed of each cells from a lognormal distribution (panel C, *µ* = 3, *σ* = 0.1). Cell movements were simulated every second and we sampled the resulting trajectories every *k* = 20 sec (panel D). The correlations observed experimentally or in simulations were evaluated using Spearman rank test (A&D) with p values from the test *ρ* = 0 being indicated on individual panels.

